# An integrated platform to systematically identify causal variants and genes for polygenic human traits

**DOI:** 10.1101/813618

**Authors:** Damien J. Downes, Ron Schwessinger, Stephanie J. Hill, Lea Nussbaum, Caroline Scott, Matthew E. Gosden, Priscila P. Hirschfeld, Jelena M. Telenius, Chris Q. Eijsbouts, Simon J. McGowan, Antony J. Cutler, Jon Kerry, Jessica L. Davies, Calliope A. Dendrou, Jamie R.J. Inshaw, Martin S.C. Larke, A. Marieke Oudelaar, Yavor Bozhilov, Andrew J. King, Richard C. Brown, Maria C. Suciu, James O.J. Davies, Philip Hublitz, Chris Fisher, Ryo Kurita, Yukio Nakamura, Gerton Lunter, Stephen Taylor, Veronica J. Buckle, John A. Todd, Douglas R. Higgs, Jim R. Hughes

**Affiliations:** MRC Molecular Haematology Unit, MRC Weatherall Institute of Molecular Medicine, Radcliffe Department of Medicine, University of Oxford, Oxford, UK; MRC WIMM Centre for Computational Biology, MRC Weatherall Institute of Molecular Medicine, Radcliffe Department of Medicine, University of Oxford, Oxford, UK; Big Data Institute, Li Ka Shing Centre for Health Information and Discovery, University of Oxford, Oxford, UK; Wellcome Centre for Human Genetics, Nuffield Department of Medicine, University of Oxford, Oxford, UK; JDRF/Wellcome Diabetes and Inflammation Laboratory, Wellcome Centre for Human Genetics, Nuffield Department of Medicine, University of Oxford, Oxford, UK; MRC Human Immunology Unit, MRC Weatherall Institute of Molecular Medicine, Radcliffe Department of Medicine, University of Oxford, Oxford, UK; WIMM Genome Engineering Facility, MRC Weatherall Institute of Molecular Medicine, Radcliffe Department of Medicine, University of Oxford, Oxford, UK; Dept. of Research and Development, Central Blood Institute, Japanese Red Cross Society, Minato-ku, Tokyo, Japan; Cell Engineering Division, RIKEN BioResource Research Center, Tsukuba, Ibaraki, Japan

**Keywords:** GWAS, gene regulation, chromatin conformation, machine learning.

## Abstract

Genome-wide association studies (GWAS) have identified over 150,000 links between common genetic variants and human traits or complex diseases. Over 80% of these associations map to polymorphisms in non-coding DNA. Therefore, the challenge is to identify disease-causing variants, the genes they affect, and the cells in which these effects occur. We have developed a platform using ATAC-seq, DNaseI footprints, NG Capture-C and machine learning to address this challenge. Applying this approach to red blood cell traits identifies a significant proportion of known causative variants and their effector genes, which we show can be validated by direct *in vivo* modelling.

## BACKGROUND

Identification of the variation of the genome that determines the risk of common chronic and infectious diseases informs on their primary causes, which leads to preventative or therapeutic approaches and insights. Whilst genome-wide association studies (GWASs) have identified thousands of chromosome regions^1^, the identification of the causal genes, variants and cell types remains a major bottleneck. This is due to three major features of the genome and its complex association with disease susceptibility. Trait-associated variants are often tightly associated, through linkage disequilibrium (LD), with tens or hundreds of other variants, mostly single-nucleotide polymorphisms (SNPs), any one or more of which could be causal; the majority (>85%) the variants identified in GWAS lie within the non-coding genome^2^. Although non-coding regions are increasingly well annotated, many variants do not correspond to known regulatory elements, and even when they do, it is rarely known which genes these elements control, and in which cell types. New technical approaches to link variants to the genes they control are rapidly improving but are often limited by their sensitivity and resolution3–6; and because so few causal variants have been unequivocally linked to the genes they affect, the mechanisms by which non-coding variants alter gene expression remain unknown in all but a few cases; and, third, the complexity of gene regulation and cell/cell interactions means that knowing when in development, in which cell type, in which activation state, and within which pathway(s) a causal variant exerts its effect is usually impossible to predict. Although significant progress is being made, currently, none of these problems has been adequately solved.

Here, we have developed an integrated platform of experimental and computational methods to prioritise likely causal variants, link them to the genes they regulate, and determine the mechanism by which they alter gene function. To illustrate the approach we have initially focussed on a single haematopoietic lineage: the development of mature red blood cells (RBC), for which all stages of lineage specification and differentiation from a haematopoietic stem cell to a RBC are known, and can be recapitulated *ex vivo* by culture of CD34^+^ progenitor and stem cells^7–9^. GWASs have identified over 550 chromosome regions associated with changes in the phenotypes of mature RBC^10, 11^; within these regions 1,114 index SNPs are in high LD with 30,694 variants, of which, only eight have been claimed as causal regulatory variants through experimental validation^12–16^.

We first identify the key cell type(s) throughout erythropoiesis by analysing enrichment of GWAS variants lying within regions of open chromatin. These regions contain the tissue-specific regulatory elements of the genome (promoters, enhancers and boundary elements). We next focus on the ∼8% of variants which lie within regulatory elements in the non-coding genome; with the remaining variants assessed for effects on coding sequences and RNA processing using established programmes^17–19^. The platform addresses the fact that both causal and non-causal variants may lie in open chromatin. Using DNaseI footprinting and a machine learning approach the platform prioritises variants predicted to directly affect the binding of transcription factors or alter chromatin accessibility^20, 21^. Having prioritised putative regulatory causal variants, the platform then links the regulatory elements in which they occur to genes using NG Capture-C, the highest resolution chromatin conformation capture (3C) method currently available for targeting numerous loci^22, 23^. To validate the predicted molecular changes caused by such GWAS variants we use CRISPR/Cas9 facilitated Homology Dependent Recombination (HDR) to directly model SNP alleles and determine their effects.

Testing our platform against 75 chromosome regions from a previous GWAS of RBC traits^11^ we identified putative causal variants at ∼80%, their candidate effector genes at ∼70%, and three or fewer candidate variants at ∼60%. By benchmarking with the eight validated causal variants from previous studies^12–16^, and genes at well characterised erythroid loci, we successfully predicted >87% of both causal variants and effector genes. Finally, we used genome editing to directly determine the *in vivo* molecular effects of candidate SNPs in two regions – showing both SNPs to be causal and verifying *JAK2* as a novel RBC trait effector gene. As this platform was developed with methods appropriate for small numbers of cells, and therefore rare cell types, the approach will enable researchers across a wide range of traits or disorders to more readily identify causal variants, the cells in which they exert their effects, their target genes, and the mechanisms by which they alter cell biology, and ultimately, disease risk.

## RESULTS

### Enrichment of variants influencing RBC traits in highly active erythroid enhancers

The first stage of an integrated platform for dissecting polygenic traits is the identification of key cell types (Supplementary Fig. 1). Recently, ATAC-seq allowed a comprehensive identification of *cis*-regulatory elements which remain constitutively present or dynamically change throughout haematopoietic lineage specification, differentiation, and maturation^8, 24^. To identify regulatory regions in early, intermediate and late erythroid cells we generated ATAC-seq from such cells obtained by *ex vivo* erythroid differentiation of CD34^+^ stem and progenitor cells^9^ from three healthy, non-anaemic individuals (Supplementary Fig. 2). We also examined ATAC-seq profiles from a variety of haematopoietic cells, including erythroid progenitors (Fig. 1a,b); in total identifying 238,918 open-chromatin regions (496-4,136 bp) present at one or more stages of erythropoiesis.

**Fig. 1.**
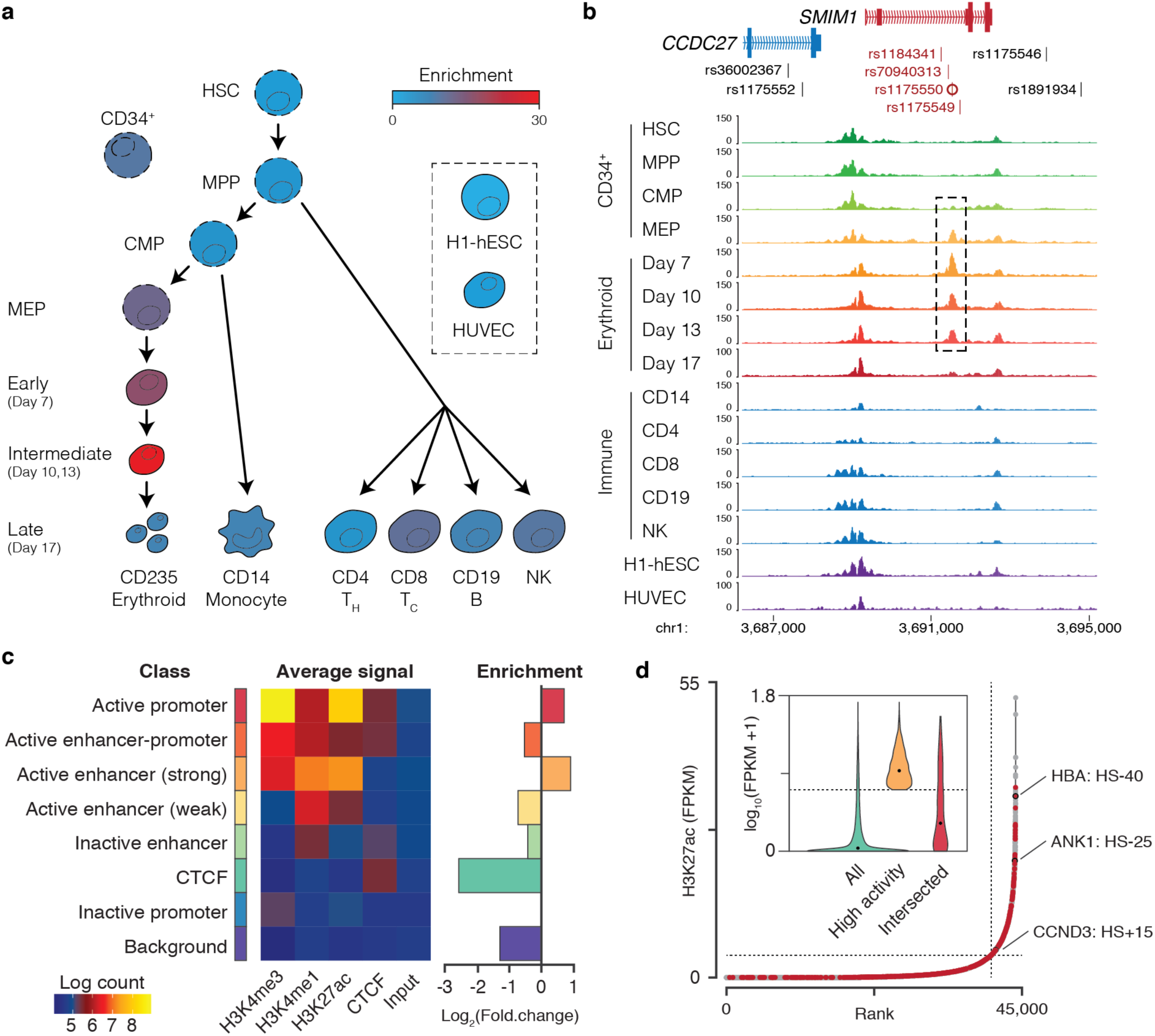
Variants associated with RBC traits lie within highly active enhancers. **a,** Schematic of selected cells from human haematopoiesis showing enrichment (-log(p) of a cumulative Binomial Distribution) for RBC associated variants within open chromatin regions of haematopoietic stem cells (HSC), multi-potent progenitors (MPP), common myeloid progenitors (CMP), megakaryocyte-erythroid progenitors (MEP), early, intermediate, and late erythroid cells from *in vitro* culture, CD14 monocytes, CD4 helper and CD8 cytotoxic T-cells, CD19+ B-cells, natural killer cells (NK), human embryonic stem cells (H1-hESC) and human umbilical vein endothelial cells (HUVEC). **b,** ATAC-seq tracks showing location of open chromatin intersecting variants (red) at the SMIM1 locus. Intersected peaks are highlighted with a dashed box. The index SNP rs1175550 is marked (circle). **c,** GenoSTAN classification and average signal of open chromatin based upon epigenetic marks with the enrichment/depletion in representation of each class amongst elements containing variants. Note, no intersection with inactive promoters was detected so was excluded from enrichment analysis **d,** Open chromatin regions distal (≥2kb) to annotated transcription start sites were ranked by level of H3K27ac ChIP-seq signal (FPKM), with highly active enhancers defined as those above the point of inflection of the curve (marked with a dashed line). Open chromatin regions containing RBC variants (dots coloured red) are enriched for highly active enhancer elements. Hypersensitive sites (HS) near important erythroid genes are shown. A violin plot of H3K27ac levels on all distal regions, highly active distal regions, and variant containing distal regions is inset; the median level is marked (black dot).

Previously, 1,114 index SNPs, each of which identifies a region of LD, have been associated with specific RBC traits in two extensive GWASs of Asian and European populations^10, 11^. These index SNPs are associated, via LD (r^2^≥0.8), with a total of 30,694 variants. Approximately 8% of these variants, covering ∼60% of RBC trait regions, intersected with open chromatin in erythropoietic cells (n=2,590). Intersections were predominantly found in fully committed, intermediate erythroid cells (days 10-13, cumulative binomial distribution, p=7×10^-29^) rather than in multipotent progenitor cells (Fig. 1a, Supplementary Figs. 3a,b). Enrichment was trait specific as variants associated with immune diseases^25–27^ showed minimal enrichment for intersection with erythroid open chromatin but strong enrichment in differentiated lymphocytes (Supplementary Fig. 3d-f) while non-haematological trait variants^28–30^ showed no enrichment in either red or white blood cells. For all traits, we saw no enrichment when we analysed ATAC-seq profiles from two non-haematopoietic cell lines.

To further characterise the intersected regulatory elements, we generated ChIP-seq data to distinguish promoters (Histone-3 Lysine-4 trimethylation, H3K4me3), enhancers (H3K4 monomethylation, H3K4me1), boundary elements (CTCF), and “active” sites (H3 Lysine-27 acetylation, H3K27ac) in committed erythroid cells. We applied GenoSTAN^31^ to assign a chromatin signature to each element and thereby generated a high-resolution map of open chromatin in erythroid cells with seven functional classes (Fig. 1c, Supplementary Fig. 4). Intersected elements were enriched for enhancers and promoters with high levels of H3K27ac but not those with low levels of H3K27ac, nor ATAC-seq peaks with CTCF enrichment (Fig. 1c). When putative enhancers were ranked for their levels of H3K27ac (Fig. 1d), elements containing RBC variants were significantly enriched amongst the highly activity erythroid enhancers (χ^2^: d.f.=3, p=7×10^-54^).

Cell-specific intersection is consistent with previous studies^32–34^, and as shown here, when applied to highly-stratified cell types may help identify the precise cells in which the variant affects function. For example, four variants including the predicted causative SNP rs1175550^13^ intersect a *cis*-regulatory element in the Small Integral Membrane Protein 1 (SMIM1; Vel Blood Group) encoding locus. This element is associated with a region of open chromatin which only appears in megakaryocytic-erythroid progenitors (MEPs) and early/intermediate erythroid precursors (Fig. 1b). A meta-analysis of all intersected open chromatin regions showed multiple trajectories of accessibility, including persistent nucleosome depletion, progenitor specific accessibility, and terminal or transient accessibility (Supplementary Figs. 5,6). While overall enrichment of predicted causal variants is strongest in intermediate erythroid cells (day 10-13), RBC traits may also be influenced by variants acting at earlier stages of erythropoiesis.

### Meta-genomic and machine learning approaches effectively prioritise causal variants

As both causal and non-causal variants fall within open chromatin, further assessment of their potential to alter the function of the underlying regulatory elements is required. We applied a combination of meta-genomics and machine learning to further characterise variants found within open chromatin in erythroid cells. Regulatory variants are likely to act by altering the dynamics of transcription factor binding, however only 10-20% of causal SNPs directly alter known transcription factor motifs^35^. This suggests that causal variants may either play an unexplained mechanistic role, or act through uncharacterised transcription factors. Sasquatch uses an unbiased approach to measure the average *in vivo* DNaseI-seq footprint for any given sequence in a specific cell type^20^, thus identifying likely transcription factor binding sites for both known and unknown transcription factors, and can therefore be used to evaluate variants in an unbiased manner (Fig. 2a). Using Sasquatch, we found 61.8% of variants in open chromatin in committed erythroid cells were predicted to have at least a weak effect on transcription factor footprints (762/1,233), accounting for variants at ∼57% of RBC LD regions (Supplementary Fig. 7). While some of these changes were found in or adjacent to known haematopoietic transcription factors, including SCL/TAL, GATA1, SPI1/PU1, NF-E2, BACH1, and MAFK, footprint changes were also seen for motifs with no known associated transcription factor (Fig. 2, Supplementary Figs. 7,8).

**Fig. 2.**
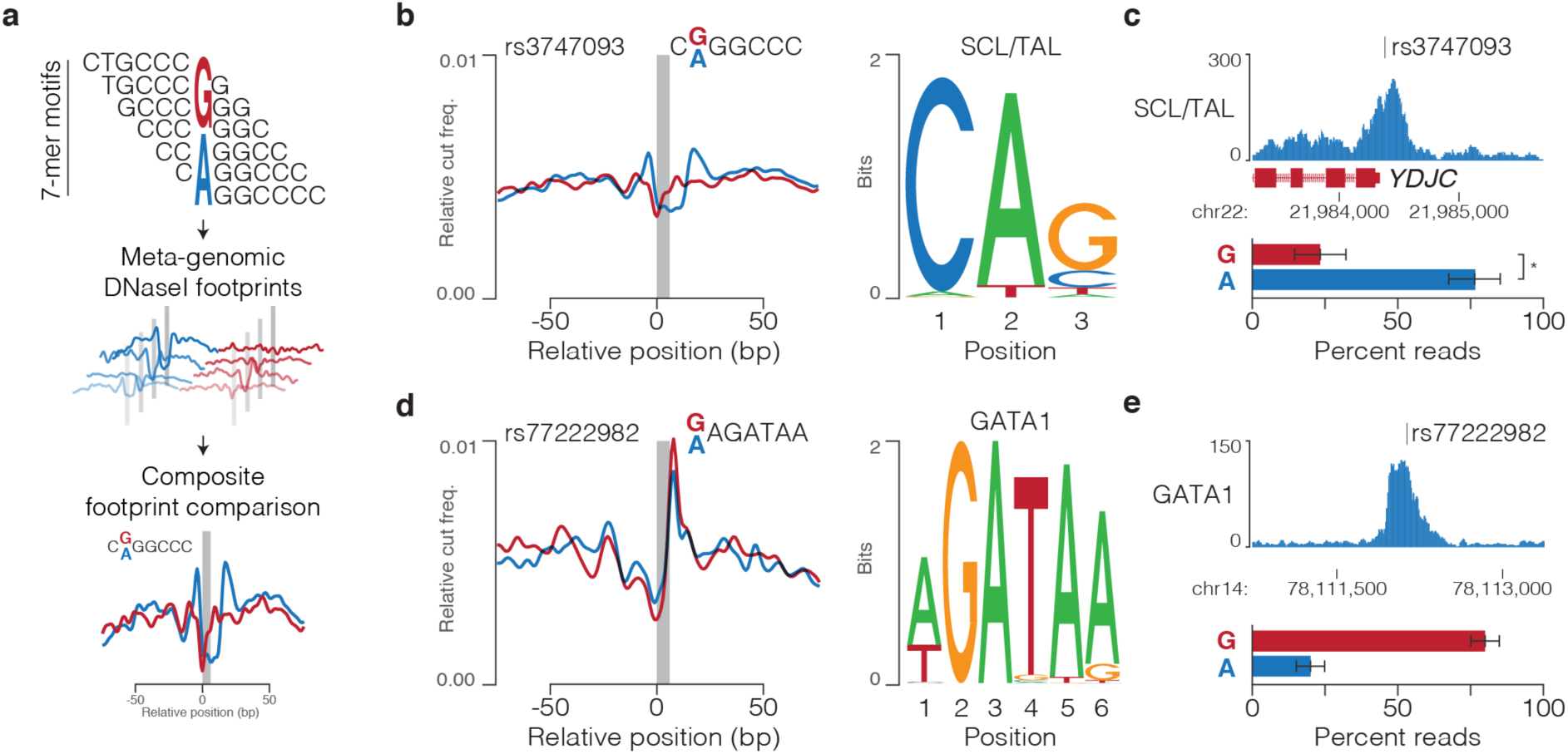
Sasquatch provides an unbiased prediction of variant effect. **a,** Sasquatch analyses *in vivo* generated DNaseI footprints over 7-mer motifs within open chromatin regions to generate meta-genomic footprints. Comparison of Relative cut frequency for each profile is used to generate predictive footprint-change scores. **b,** rs3747093 is within a 7-mer motif (grey bar) which is predicted to alter the DNaseI footprint of SCL/TAL based on presence of the SCL/TAL binding motif. **c,** SCL/TAL ChIP-seq shows allelic skew over rs3747093 as shown by percent of reads containing either allele (*P=0.0468, Ratio paired t-test, n =3). **d,** rs77222982 is within a 7-mer motif (grey bar) which is predicted to alter the DNaseI footprint of GATA1 based on presence of the GATA1 binding motif. **e,** GATA1 ChIP-seq shows allelic skew over rs77222982 as shown by percent of reads containing either allele (n=2). Error bars depict standard error of the mean.

Changes in specific transcription factor binding predicted by Sasquatch were validated by analysis of heterozygous variants using ChIP-seq. Notably, for rs3747093 which falls within an SCL/TAL binding motif, significant allelic imbalance was seen in erythroid SCL/TAL ChIP-seq in three independent datasets (Fig. 2b-c). Similarly, rs77222982 which is directly adjacent to an AGATAA motif showed allelic imbalance in GATA1 binding (Fig. 2d-e). Often, skew in enrichment was seen across more than one factor in elements affected by a single variant, probably reflecting co-dependency in their binding (Supplementary Fig. 8). Such imbalance is consistent with the alteration of binding predicted by Sasquatch, demonstrating its ability to accurately detect causative variants.

Convolutional neural network based machine learning can predict open chromatin^36, 37^ and was also used to identify causal variants. We used 936 chromatin-accessibility and epigenetic datasets to train a deep convolutional neural network, deepHaem^21^, to predict chromatin accessibility based on DNA sequence across haematological cell-types (Fig. 3a, Supplementary Fig. 9a-d). Using deepHaem it is possible to predict the effect of variants on chromatin accessibility. DeepHaem identified 91 variants in open chromatin with changes greater than 10% of the maximum accessibility score (1.0), with the strongest effects seen in MEP and erythroid populations (Fig. 3b). 45 of the variants predicted to alter chromatin-accessibility had scores greater than 0.1 in erythroid cells. Using ATAC-seq, we identified heterozygous alleles for 15 of these 45 variants in three healthy individuals. Comparison of sequencing from these alleles showed significant bias in ATAC-seq accessibility at seven of the sites and skew at a further five sites (Fig. 3c, Supplementary Fig. 9e) indicating that deepHaem can accurately predict variant-induced changes in chromatin accessibility.

**Fig. 3.**
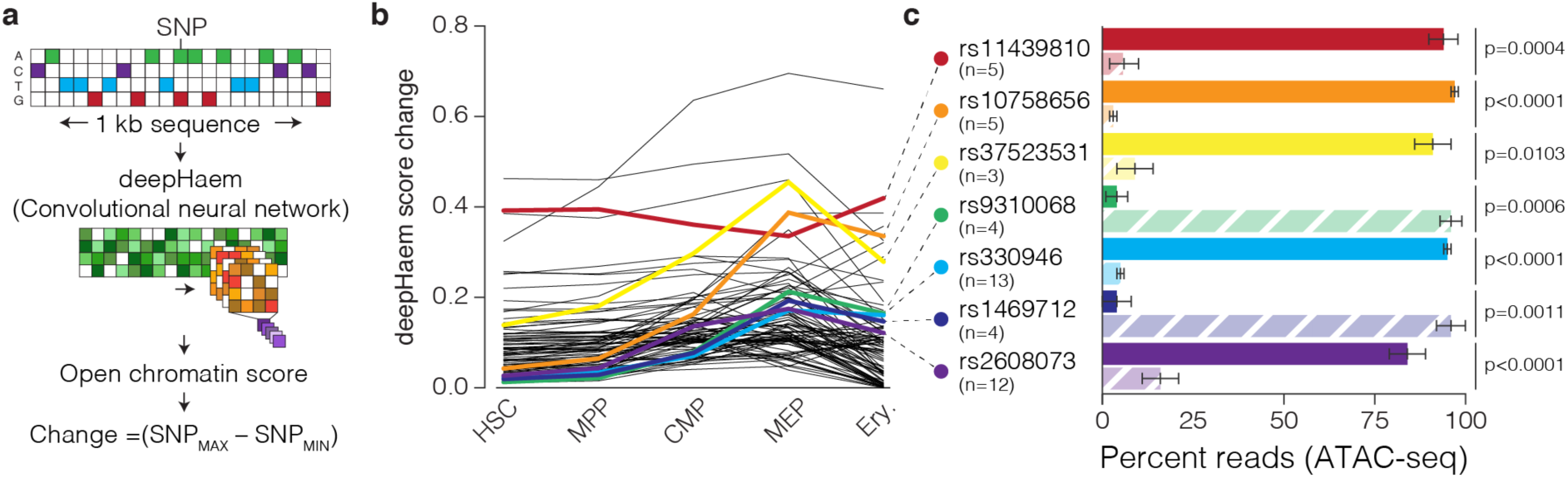
Deep Learning predicts variant driven changes in chromatin accessibility. **a,** deepHaem, a deep convolutional neural network, calculates a chromatin openness score using 1 kb of DNA sequence which can be used to compare variant alleles. **b**, Comparison of alleles for all RBC trait variants in open chromatin (n=2,662) identifies variants with a predicted to change deepHaem openness scores by more than 0.1, or 10% of the maximum openness score (n=91). **c,** Mean percentage of day 10 and day 13 erythroid ATAC-seq reads on either the reference (dark bar) or variant (light dashed bar) allele from heterozygous individuals with a minimum of 5 reads. Error bars depict the standard error of the mean with the number of independent replicates from either multiple donors and/or multiple differentiations shown in parentheses. p-values shown are for a ratio paired t-test.

To assess how well the platform performed at identifying causative regulatory variants we used previously characterised RBC trait variants. Currently, no RBC trait variants have been definitively shown to be causative using direct *in vivo* modelling; however, eight regulatory variants have strong support from functional assays^12–16^. The approach established here identified that seven of these eight variants lie in open chromatin in erythroid cells, and therefore had the potential to be regulatory causal variants. Characterisation with Sasquatch or deepHaem further prioritised six of these variants as likely to be causative (Supplementary Table 1). Therefore, the platform accurately prioritises causal variants, and thus identifies variants for functional analysis.

### A comprehensive search for all causal variants within an RBC GWAS

The first major RBC trait GWAS identified 75 index SNPs^11^; the associations identified are likely to represent the most common variants with moderate effect sizes and some rare variants with large effect sizes, therefore we focused specifically on this dataset for in-depth follow-up. A comprehensive GWAS decoding platform must prioritise causal variants by treating all mechanisms as plausible. By examination of the 75 index SNPs, as well as variants in high LD with them (r^2^≥0.8, 1000 Genomes Project; n=6,420), we identified 486 candidate regulatory variants within 61 of the 75 chromosome regions (Supplementary Table 1). In addition to regulatory variants, we considered the possibility for coding and splicing changes across these regions. Putative coding sequence changes were identified using ANNOVAR^17^ and then filtered for erythroid expressed genes. This identified 20 variants predicted to alter protein sequence at 14 regions (Supplementary Table 1). Next, putative alternative-splicing variants were identified using a combination of Splicing Index^18^ and a deep learning approach, SpliceAI^19^. Together, these programmes identified 13 putative splice-altering SNPs in 11 erythroid expressed genes across nine regions; however, no variants were highlighted by both algorithms (Supplementary Table 1). Using these integrated analyses for coding, splicing and regulatory mechanisms we identified candidate causal variants at 63 of the 75 chromosome regions, with 43 of these having three or fewer strong candidates, and the majority of candidates being in tight linkage (r^2^≥0.9; n=394/515) with their index SNP (Fig. 4, Supplementary Figs. 10, 11a).

**Fig. 4.**
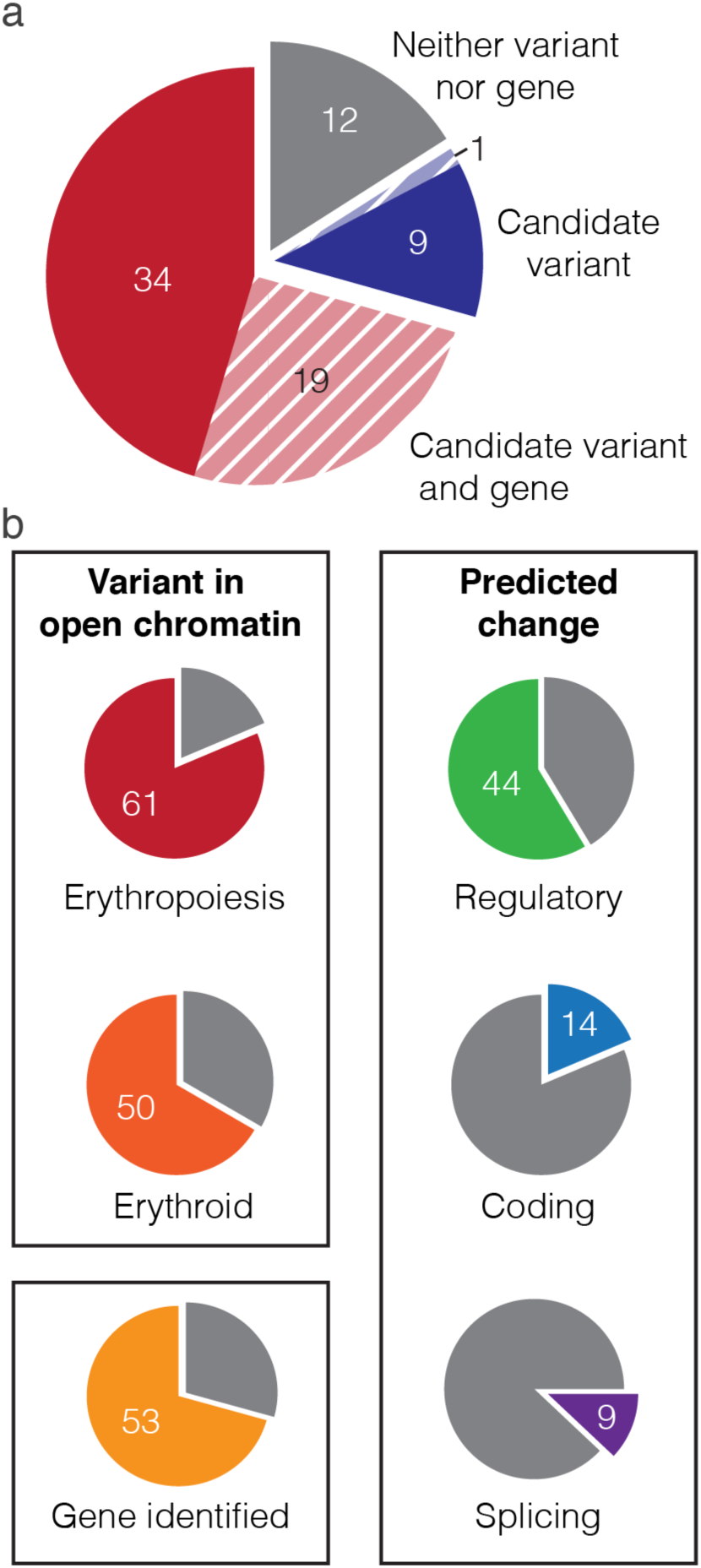
The integrated experimental and bioinformatics platform identifies candidate causal variants and effector genes at the majority of polygenic trait regions. **a,** Cumulative analysis of RBC trait associated variants at 75 GWAS chromosome regions identified candidate causal variants in 63 regions with three or fewer candidate causal variants at 43 regions (solid colouring) and more than three candidates at 20 regions (pale striped colouring). **b,** Pie charts with the number of regions, from a total of 75, with variants found in open chromatin, with variants predicted to alter a regulatory site (Sasquatch, deepHaem), or coding sequence (ANNOVAR), or splicing (SpiDEX, SpliceAI), and regions with identified candidate effector genes. 16 regions have candidate causal variants with separate mechanisms of action, five of which have regulatory, splicing and coding candidates.

We next considered why no causal variants were predicted for 12 chromosome regions. Immediate possibilities are that the variant affects gene function in a way that is currently unrecognised, or exerts effects in an untested cell type. Consistently, rs855791, also identified in multiple GWASs, including for iron status^38, 39^, is a missense variant of *TMPRSS6*. *TMPRSS6* is expressed in the liver and encodes Matripase-2, a suppressor of the iron homeostasis master regulator, Hepcidin^40, 41^. It is also possible that the causal variant affects mRNA stability. However, there are currently no good predictive software programmes for this. Finally, it may be that causal variants were not identified because the initial GWAS study was not sufficiently powered or used a sub-optimal catalogue of variants; resulting in incompletely resolved genetics. Indeed, index SNPs in unresolved loci were less likely to be replicated in subsequent RBC trait GWASs^10, 42^ than index SNPs at resolved loci (Supplementary Fig. 11b). Additionally, in a region with multiple unlinked causal variants, incompletely resolved genetics can lead to index SNPs being identified through weak association (r^2^<0.8) with two or more causal variants. Such index SNPs are referred to as tag SNPs^43^ (Supplementary Fig. 11a). At the *TMCC2* locus, where rs9660992 is an index SNP^11^, moderate linkage (r^2^=0.51-0.82) is seen with two independent index SNPs from a subsequent RBC trait GWAS^10^. While rs12137294 and rs1172129 are themselves unlinked (r^2^=0.46), each is in tight linkage with several variants in open chromatin (Supplementary Fig. 12); suggesting rs9660992 may be a tag SNP. Therefore, both additional cell types and incomplete genetics can explain unresolved regions.

### High resolution 3C mapping accurately identifies effector genes

The target or effector genes for splicing and coding variants can be directly inferred, but the effector genes of regulatory variants must be identified experimentally. For enhancers to regulate gene expression they often physically interact with target promoters, likely through loop-extrusion and/or phase-separation^44, 45^. The close proximity required for regulation can be identified by chromosome conformation capture (3C) to map interactions^46^ and this provides a means by which to identify effector genes. NG Capture-C uses biotinylated oligonucleotide probes to target specific loci at high resolution in multiplexed 3C samples^23^; allowing statistical comparison for identification of enhancer-promoter interactions. We designed probes for 214 variant containing *cis*-regulatory elements covering 53/61 chromosome regions with putative regulatory variants; then simultaneously generated 3C interaction data in intermediate erythroid cells, H1-hESCs and HUVECs to link *cis*-regulatory elements with their effector genes. Using a combination of tissue-specificity (DESeq2)^23^, Bayesian modelling (Peaky)^47^, and promoter proximity (≤5 kb) to call variant-promoter interactions we identified 194 candidate effector genes at 48 of the 53 targeted regions (Fig. 5a-b, Supplementary Table 1). For each targeted region, NG Capture-C identified an average of four genes, which is consistent with the predicted number of gene targets for enhancers^48–50^, though whether multiple genes contribute to a GWAS trait at a single locus remains to be determined. Although some methods have indicated that GWAS variants are most frequently found within 20 kb of their target genes^51^ we detected interactions up to 992 kb, with a median distance of 83.9 kb (±9.3 kb SE, Fig. 5c). Such long-range interactions were seen at several well characterised erythroid loci including *CITED2* (139 kb), *SLC4A1* (47 kb), *RBM38* (24 kb), *ANK1* (25 kb), *MYB* (85 kb), and *HBA1/2* (63 kb), showing that GWAS variants, as for other enhancer-promoter interactions, may act over large distances (Fig. 5d, Supplementary Figs. 13-18).

**Fig. 5.**
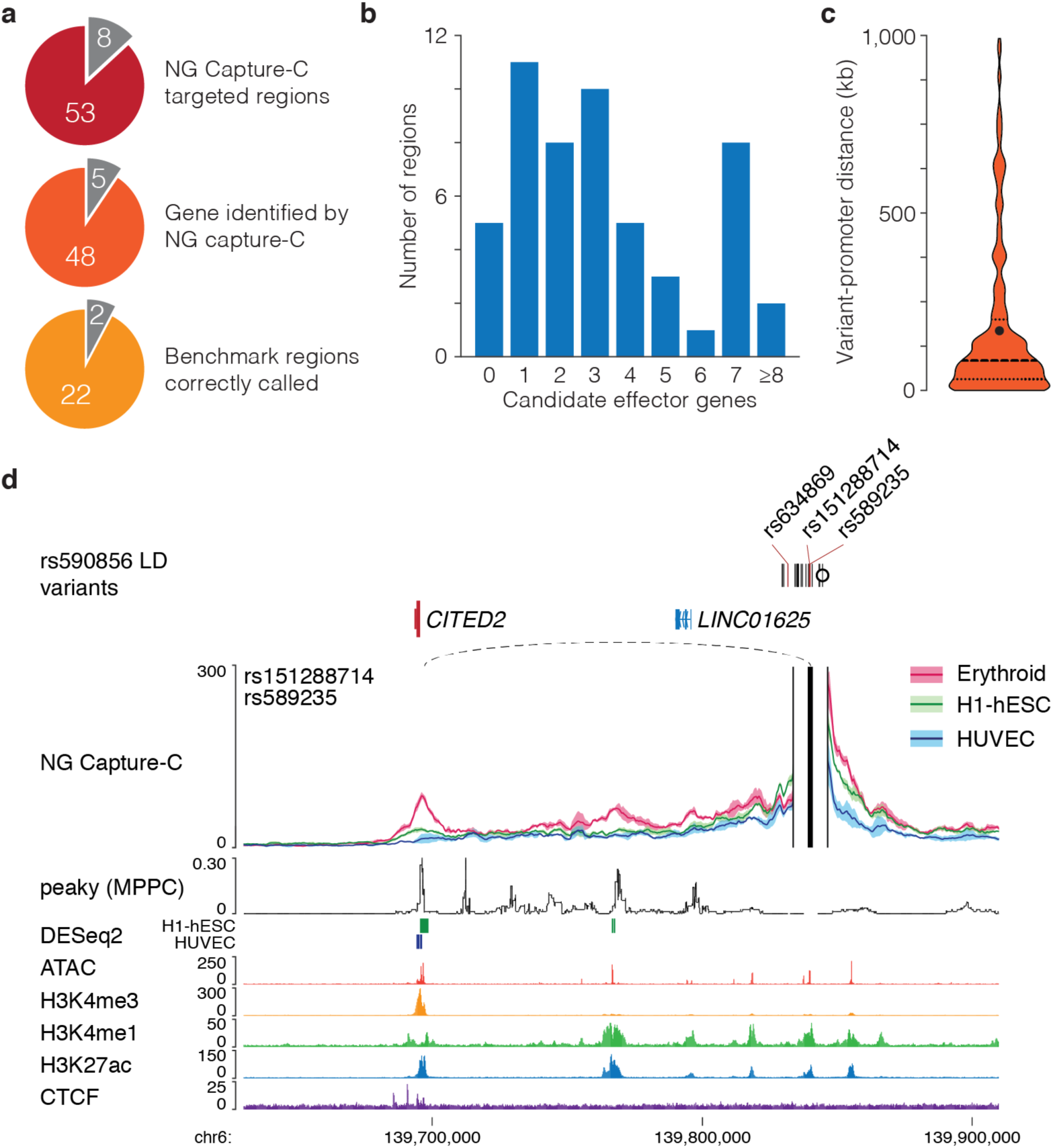
NG Capture-C can detect long range variant-promoter interactions. **a,** NG Capture-C oligonucleotides were designed for 61 chromosome regions with candidate regulatory causal variants, regions were excluded from targeting based on sheer number of targets (>100 target sites at a single locus, n=1), or where probes were impossible to design due to repetitive elements. Following analysis of generated 3C data genes were identified at 48 of the 53 targeted regions, 24 of which were used for accuracy benchmarking. **b,** Histogram of the number of candidate effector genes identified by NG Capture-C at each region. **c,** Violin plot of the distance between variants and the target transcription start sites. Median (83,944 bp) shown as a thick dashed line and mean (168,148 bp) shown as a black circle. **d,** 3C interaction profile for open chromatin containing rs151288714 and rs589235 in erythroid, human embryonic stem (H1-hESC) and human umbilical vein endothelial (HUVEC) cells (n=3). Capture viewpoints and proximity exclusion regions (solid vertical lines) were designed for open chromatin regions and profiles show mean interactions (solid line) with one standard deviation (shading). *CITED2*-variant interactions were identified as erythroid specific interactions (dashed loops; DESeq2 q-value < 0.05 shown as bars). Peaky values depict the Marginal Posterior Probability of Contact (MPPC) in erythroid cells. Variants within open chromatin are red, as are variant interacting genes, the index SNP is marked with a circle. FPKM normalised ATAC-seq and ChIP-seq tracks are from erythroid cells. Interaction was found with *CITED2*, which encodes the Cbp/p300 Interacting Transactivator with Glu/Asp (E/D)-rich tail 2 protein and required for normal haematopoiesis.

The erythroid system and RBC traits have been intensively analysed and characterised; therefore, we were able generate a set of the 24 “most likely” effector genes within the 53 targeted chromosome regions based on prior knowledge of their function (Supplementary Fig. 19a). This set of genes allowed us to benchmark our approach; finding that with NG Capture-C we correctly identified 22 of the 24 most-likely effector genes (Fig. 5a, Supplementary Fig 19b). Of the remaining regions, no genes were identified at one (*miR-181a*), and four incorrect candidates were identified in the region where *TAL1* is almost certainly the effector. With these exceptions, NG Capture-C performs with a high rate of success in identifying effector genes linked to potential causal variants. Three previous attempts with diverse methods to identify effector genes associated with RBC traits have been reported^5,6, 11^. These were an annotation-based approach^11^, Promoter Capture-HiC^5^ (PC-HiC), and a gene-centric shRNA screen^6^. We directly compared these different approaches with NG Capture-C. There was little consistency between the candidate gene lists from these approaches (Supplementary Fig. 19c), with NG Capture-C the only method to identify *HBA-1/2*, the α-globin encoding genes, which are known to be associated with anaemia and changes in RBC traits^52^ (Supplementary Figs. 17, 19c). Our approach was also unique in identifying *RPL19*, of interest because ribosomal genes are known to cause Diamond-Blackfan anaemia^53^. Across the 24 benchmark regions, NG Capture-C and the annotation-based approach were the most sensitive methods, respectively identifying 91.7% and 70.8% of the most-likely effector genes correctly (Supplementary Fig. 19 b,d). Overall, the direct comparison of different gene identification methods shows that NG Capture-C is the most successful tool.

### Direct modelling of rs9349205 shows reduced expression of its target gene *CCND3*

The most direct evidence that a particular variant alters gene expression comes from introducing both alleles to an isogenic background and observing an appropriate change in the relevant cell type. Previous studies characterising RBC trait variants have used reporter assays and/or targeted deletions of the regulatory element^13–16^. However, these may not faithfully recapitulate variants effects *in vivo.* Therefore, we used CRISPR/Cas9-facilitated homology directed repair (HDR) to directly model prioritised variants at five GWAS regions in the Human Umbilical Derived Erythroid Progenitor (HUDEP-2) cell line (Supplementary Fig. 20, 21); a model of human erythroid differentiation and maturation^54, 55^. As previous studies have shown some clonal variation when using such cells^56^ it is essential to analyse multiple independently isolated clones. We were able to generate sufficient independent clones for robust analysis of two regions (*CCND3* and *JAK2*).

Using our platform, rs9349205 was identified as tightly linked to the index SNP (rs9349204, r^2^=0.841); rs9349205 is the only one of ten linked variants which lies within open chromatin in committed erythroid cells, and shows 3C interaction with *CCND3* (Fig. 6a), which is the most-likely effector gene^16^. rs9349205 also had small effects on both the Sasquatch DNaseI footprint and deepHaem chromatin openness scores (Supplementary Figs. 20,22a). Editing was used to convert rs9349205^A/A^ HUDEP-2 cells to rs9349205^G/G^; a non-erythroid locus was also edited to control for non-specific effects from editing (e.g. spontaneous differentiation). Using ATAC-seq to assess chromatin accessibility, we found that the identified regulatory element in the rs9349205^G/G^ genotype was 54.5% less accessible than in rs9349205^A/A^ cells (Fig. 6b, Supplementary Fig. 22b). The rs9349205^G/G^ clones also showed significantly reduced *CCND3* expression during erythroid differentiation (Fig. 6c). As previously discussed^16^, *CCND3* regulates the G2 to S transition during erythropoiesis, and thus knockout of *CCND3* in mice leads to an increased erythrocyte volume, consistent with linkage to changes in mean cell volume (MCV) detected through GWAS.

**Fig. 6.**
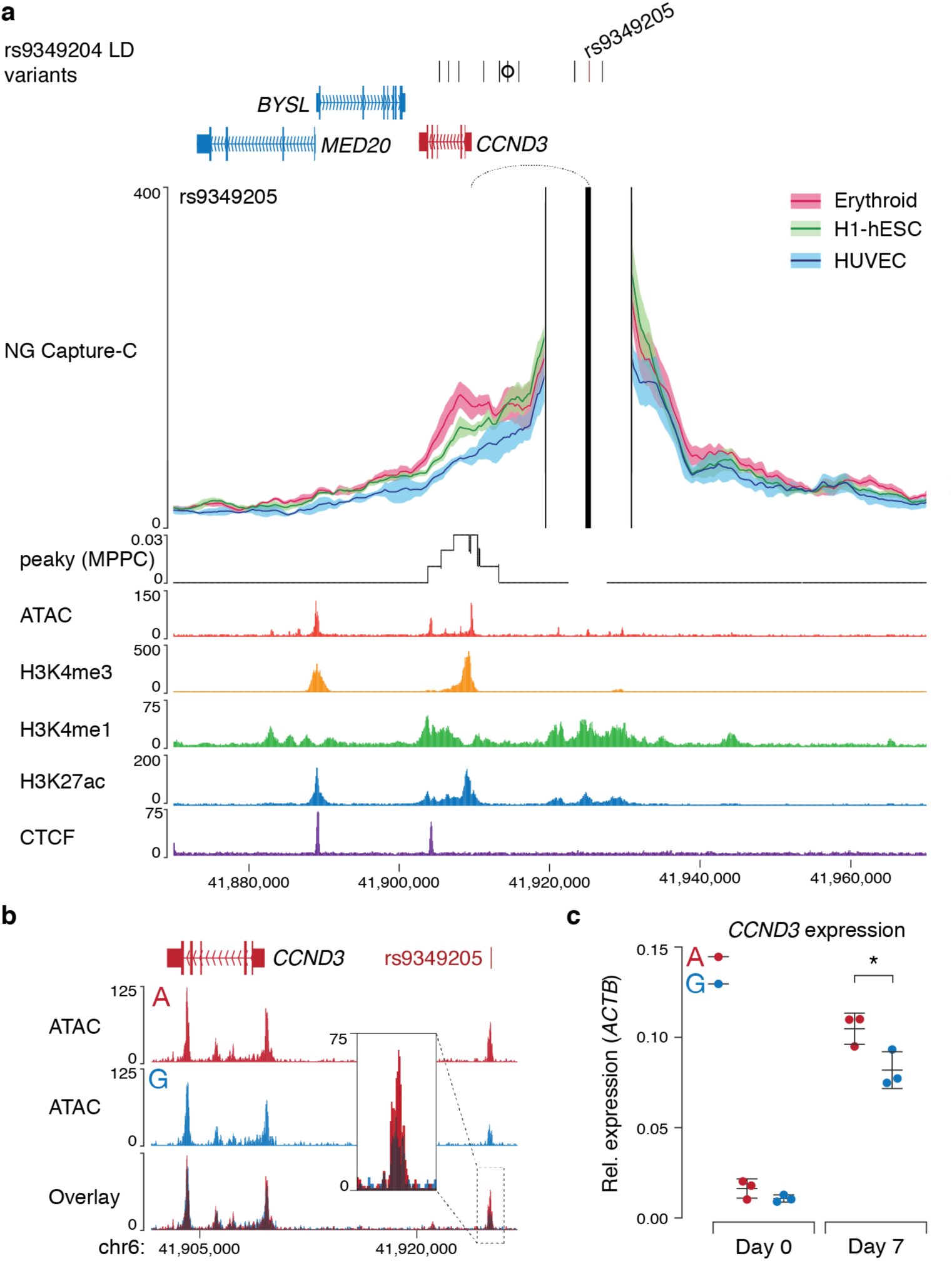
rs9349205 interacts with, and regulates CCND3. **a,** 3C interaction profile for rs9349205 in erythroid, embryonic stem (H1-hESC) and umbilical vein endothelial (HUVEC) cells (n=3). Profiles show windowed mean interactions (solid lines) with one standard deviation (shading). Peaky values depict the MPPC in erythroid cells. Interaction with *CCND3* was detected by peaky (dotted loop; MPPC > 0.01). Variants within open chromatin are red, as are variant interacting genes, the index SNP is marked with a circle. FPKM normalised ATAC-seq and ChIP-seq tracks are from erythroid cells. **b,** Merged FPKM normalised ATAC-seq (n=3) from HUDEP-2 cells homozygous for either rs9349205 allele with overlaid track showing high similarity, and a slight reduction at the intersected peak for homozygous G clones (inset). **c,** Real time reverse-transcriptase PCR of *CCND3* in differentiating HUDEP-2 clones (n=3) showed lower expression in G clones at day 7 (Student’s two-tailed t-test, *p=0.0387). Bars show mean and one standard deviation of independent clonal populations (circles).

### rs10758656 causes reductions in chromatin accessibility and *JAK2* expression

Using NG Capture-C we identified *JAK2*, which encodes Janus Kinase 2, as an effector gene for variants in high LD with the index SNP rs2236496. We confirmed this interaction using NG Capture-C from the *JAK2* promoter (Fig. 7a). Of the 18 linked variants, only two, rs10758656 and rs10739069, intersect open chromatin. Of these two SNPs Sasquatch characterised rs10758656 but not rs10739069 as having the potential to affect transcription factor binding, with the motif strongly matching that of the GATA1 binding motif (Fig. 7b, Supplementary Fig. 23a). DeepHaem also predicted that rs10758656 but not rs10739069 would affect chromatin accessibility. Therefore, HUDEP-2 cells, which are heterozygous A/G for rs10758656, were edited to homozygosity. We generated 16 independent clones homozygous for either A (n=10) or G (n=6). ATAC-seq of these cells during expansion and differentiation showed up to 82% ablation of open chromatin in the rs10758656^G/G^ clones, associated with 86.2% and 58.4% reductions in GATA1 binding and H3K27ac, respectively (Fig. 8a-b, Supplementary Figs. 24b-e). These findings match the prediction of both Sasquatch and deepHaem. Despite being closer to the promoter of *RCL1* than *JAK2*, rs10758656^G/G^ specifically reduced expression of *JAK2* (Fig. 8c, Supplementary Fig. 24f,g), consistent with the specificity of the rs10758656-*JAK2* interaction profile seen in NG Capture-C. JAK2 functions as part of the erythropoietin signalling pathway^57^. Our results demonstrate that *JAK2* is a GWAS effector gene and most likely results in changes to the MCV noted in GWAS through altered signalling responses.

**Fig. 7.**
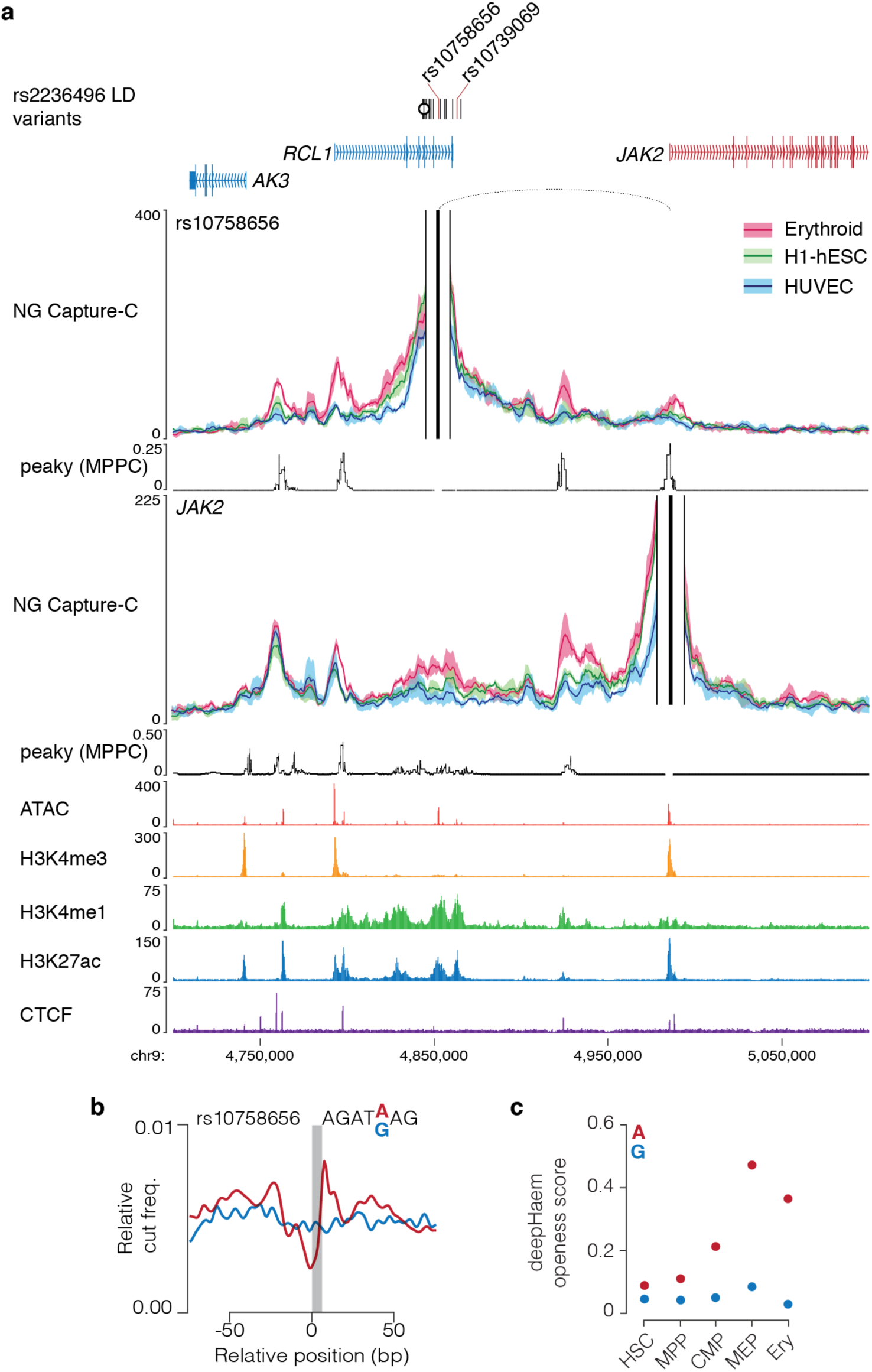
rs10758656 interacts with *JAK2* and is predicted to alter chromatin accessibility. **a,** 3C interaction profiles for rs10758656 and *JAK2* in erythroid, embryonic stem (H1-hESC) and umbilical vein endothelial (HUVEC) cells (n=3). Profiles show windowed mean interactions (solid lines) with one standard deviation (shading). Peaky values depict the MPPC in erythroid cells. *JAK2*-rs10758656 interaction was detected by peaky (dotted loop; MPPC > 0.01). Variants within open chromatin are red, as are variant interacting genes, the index SNP is marked with a circle. FPKM normalised ATAC-seq and ChIP-seq tracks are from erythroid cells. **b,** Sasquatch profiles for rs10758656 show loss of a GATA footprint. **c,** deepHaem openness scores rs10758656 predict a loss of chromatin accessibility in erythroid cells.

**Fig. 8.**
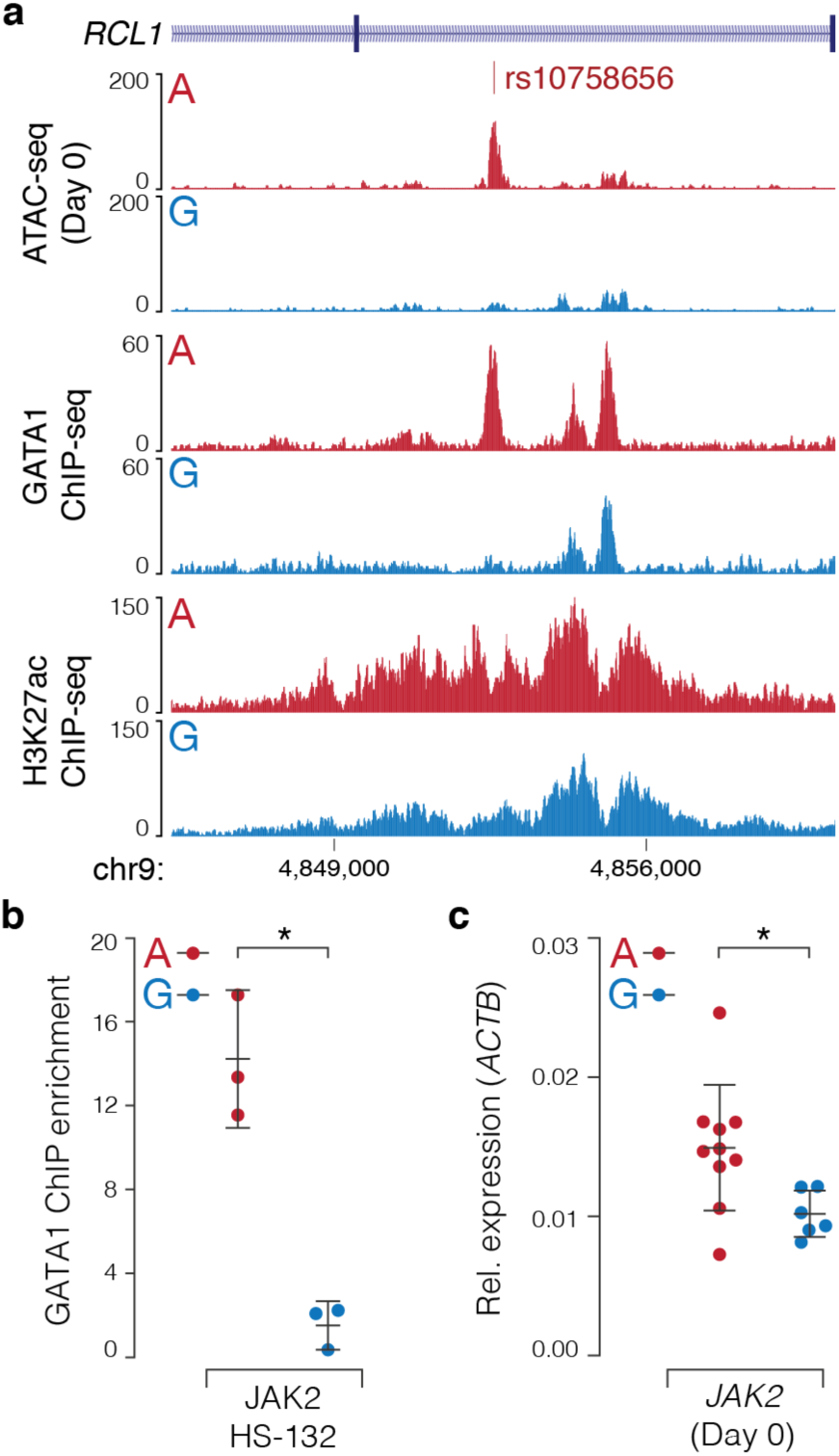
rs10758656 causes loss of open chromatin and reduced *JAK2* expression. **a,** FPKM normalised ATAC-seq (n=3), GATA1 ChIP-seq (n=3) and H3K27ac ChIP-seq (n≥1) from differentiating HUDEP-2 clones homozygous for either rs10758656 allele. **b,** Real time quantitative PCR for GATA1 ChIP at rs10758656 (*Student’s two-tailed t-test, p=0.0136). **c,** Real time reverse-transcriptase PCR of *JAK2* in differentiating HUDEP-2 clones (n≥6) showed lower expression in G clones (Mann-Witney test, *p=0.0160). Each circle reflects an independently isolated clonal population, bars show mean and one standard deviation.

## DISCUSSION

Here we have developed and validated a platform to identify causative GWAS variants and link them to the genes whose function they affect. In our platform ATAC-seq analysis allows researchers to identify relevant cell types using the fundamental regulatory elements of the genome: enhancers, promoters and boundary elements. GWAS variants are then assessed *in silico* to predict which variants are likely to alter gene expression or function. Candidate regulatory variants are finally linked to their effector genes using NG Capture-C. Using this method to analyse variants in high LD to RBC trait index SNPs resulted in identification of candidate causal variants and effector genes at a majority of chromosome regions (>70%). Benchmarking also shows that this approach is robust, with 88% of validated causal variants, and 92% of most-likely effector genes identified. Application of this method to fine-mapped GWAS variants is likely to further improve its success. Finally, the functional effects of candidate polymorphisms can then be assessed using allele-specific assays of chromatin accessibility and gene expression.

This platform has been developed and benchmarked using data from purified haematopoietic cells at various stages of commitment, differentiation and maturation along the erythroid pathway to producing RBC. Using haematopoiesis as a model, we show how causal variants can be assigned to the cell types in which they exert their effects and the genes whose expression is perturbed. In principle, this method could be used for any GWAS datasets for which appropriate cell types are available. To ensure that this would apply to rare cell types and a wide range of diseases, we have established a platform that can be effectively applied using as few as 500 cells for ATAC-seq and 20,000 cells for NG Capture-C^24, 58^. These data can then be used to improve *in silico* processing and machine learning, meaning that damaging changes can be predicted and prioritised using data from rare cells and those grown under varying conditions of stimulation.

Linking variants to their effector genes using NG Capture-C can easily and reproducibly be applied across a wide range of cell-types at hundreds of specifically targeted sites in either gene- or enhancer-centric designs^22, 23^. The ability to compare 3C data from multiple cell types allows tissue-specific and tissue-invariant interactions to be called by a wide range of statistical approaches^23, 47, 59–61^, increasing the throughput of accurate effector gene identification. These candidates can then be validated with functional follow-up, such as screening approaches^6^, or as shown here, *in vivo* modelling, to help to explain associated cellular phenotypes.

In addition to elucidating the genes involved at GWAS regions, we have also addressed the multiple molecular mechanisms that may underlie such signals. To date, strong evidence supports a mixture of coding, splicing and regulatory mechanisms^62^. The approach described here identifies enhancer, promoter, RNA processing and coding variants. Despite this we were still unable to identify causative variants at 16% of chromosome regions. This could partly have resulted from the fact that initial variant identification used linkage disequilibrium, which could be improved with either fine mapping or whole genome sequencing^62, 63^. Nevertheless, other factors are also likely to contribute. The cell types affected in any complex disease are not necessarily the most obvious candidates. For RBC traits, the most likely affected lineage is erythropoiesis itself. However, other cell types modify erythropoiesis, including those producing growth factors, cytokines, or mediating cell/cell interactions such as macrophages that facilitate enucleation of RBC precursors. Furthermore, causal variants may act in the identified cell type, but only in response to specific environmental cues or signalling. Therefore, platforms such as this must be implemented with a comprehensive appreciation of the systems involved. It is also important to consider that additional untested molecular mechanisms may underlie GWAS signals. Although we found no specific enrichment for variants in CTCF elements, numerous were within CTCF peaks. Recent evidence has shown that disruption of CTCF binding by common variants plays a role in determining the severity of influenza and breast cancer^64, 65^, thus it likely represents a less common, yet important molecular mechanism. Additionally, modelling of rs10758656 showed near complete loss of open chromatin. It is equally possible that some variants generate, rather than abolish, open chromatin sites. Such a variant has already been described as causing anaemia at the α-globin locus^66^. These sites could only be detected by analysis of individuals with the correct genotype.

## CONCLUSIONS

Although this integrative platform efficiently identifies variants and genes, it shifts the bottleneck of GWAS follow-up to validation. Using direct *in vivo* modelling we have shown how alleles can alter enhancers and gene expression to different extents. Although we have shown in principle that prioritised variants can be proven to be functionally causative it requires an HDR editable cell type, is labour intensive, and does not work at all loci; this step will require rapid single base editing to enable significant progress. We expect that with the implementation of integrative platforms such as this, and with ongoing advancement of molecular techniques and editing technologies the benefits of GWAS for understanding human physiology and improving health will accelerate.

## METHODS

### Separation of blood cells

Fresh blood was sourced either as whole blood collected from three healthy donors (two males, one female) using EDTA Vacuettes (Becton Dickson) or 5 ml leukocyte cones (NHS Blood & Transport). Whole cell counts were performed on a Pentra ES60 (Horiba) for donor blood to ensure clinically healthy red blood cell counts (Supplementary Fig. 2). Blood was diluted with PBS and overlaid onto Histopaque-1077 (Sigma) and centrifuged for 30 min at 630 rcf (no brake). Peripheral Blood Mononuclear Cells (PBMCs) were washed in PBS and MACS buffer (PBS, 2 µM EDTA, 0.5% BSA) and stained with Human CD34 Microbead kit (Miltenyi Biotec) following the manufacturer’s instruction for 30 minutes (4 °C) before being passed successively through two LS Columns (Miltenyi Biotec) with three MACs buffer washes. Counting of cells was performed on a Luna FL (Logos) after staining with acridine orange (AO) and propidium iodide (PI). CD34^+^ cells were either stored in freezing buffer (90% FBS, 10% DMSO) or resuspended in Phase I medium for a three-phase differentiation^9^. CD34^+^ depleted PBMCs were then sequentially stained and passed over LS or MS columns for selective purification of CD8+, CD14+, CD4+, CD19+ and, NK cell populations using cell-type specific kits (Miltenyi Biotec). For NK cells, non-NK cells were first blocked with a biotin-antibody cocktail before binding to NK microbeads following the manufactures instructions.

### Differentiation of CD34^+^ cells

Cells were differentiated under a three phase *ex vivo* protocol adapted from that used for the BEL-A cell line^7, 9, 67^. Growth media are listed in Supplementary Table 2. Briefly, for differentiation 0.5-2.5×10^5^ cells were resuspended on day 0 in Phase I media at 10^5^ cells ml^-1^. Cell counts were performed on days 3 and 5 with additional Phase I media added to return the concentration to 10^5^ cells ml^-1^. On day 7, cells were counted and pelleted (400 rcf, 5 min, RT) and resuspended in Phase II media at 3×10^5^ cells ml^-1^. Cells were counted on day 9 and diluted to 3×10^5^ cells ml^-1^ Phase II media. On day 11, cells were counted and pelleted (400 rcf, 5 min, RT) and resuspended in Phase III media at 10^6^ cells ml^-1^. Cells were counted on days 13 and 15 and diluted to 10^6^ cells ml^-1^ in Phase III media. Reproducibility between differentiations was confirmed morphologically with cytospins, immunologically with six FACS cell surface markers and epigenetically with ATAC-seq enhancer staging^24^. For morphological analyses 10^5^ cells were resuspended in 200 ml PBS and spun (5 min, 400 rpm) in a Cytospin 4 (ThermoFisher), before staining with modified Wright’s Stain on a Hematek (Bayer Health Care), and mounting with DPX (Sigma). Images were taken on an Olympus BX 60 microscope at 10x and 20x magnification. For differentiation and enucleation FACS analyses 10^5^ cells were resuspended in FACS buffer (90 % PBS, 10 % FBS) and stained with an erythroid differentiation panel of antibodies (Supplementary Table 4) against CD34, CD36/Fatty acid translocase, CD235a/Glycophorin A, CD71/Transferrin Receptor, CD233/Band3, CD49d/α-Integrin, and with Hoescht-33258 (ThermoFisher) for live/dead analysis, with Hoescht-33342 (ThermoFisher) for enucleation assays. For immune cell purities, cells were stained with cell-type specific panels of antibodies (Supplementary Fig. 25, Supplementary Table 4). FACS was carried out on an Attune NxT (ThermoFisher), voltages and compensation were set using Ultra Comp eBeads (ThermoFisher) for antibodies, and single stained cells for dyes. Gating was performed using fluorescence minus one (FMO) controls. Analysis was performed using either Attune NxT software (v3.0) or FlowJo (v10.4.2).

### Cell line culture and HUDEP-2 differentiation

Human ESC line H1 (H1-hESC; WiCell) was grown on Matrigel (Corning) coated plates in mTeSR1 medium (StemCell technologies). Cells were harvested as a single cell suspension using Accutase (EDM Millipore); ATAC-seq and fixation were carried out in mTeSR1 medium. Primary neonatal Human Umbilical Vein Endothelial Cells (HUVEC) were sourced from three suppliers to provide genetic diversity (Lonza, Gibco, PromoCell). HUVECs were expanded in endothelial cell growth medium (Sigma) up to five passages following the manufacturer’s protocol. Briefly, HUVECs were grown to 60% confluence, washed with HBSS at room temperature and sub-cultured following light trypsination using Trypsin-EDTA (Sigma) at room temperature and terminating the reaction with trypsin inhibitor (Sigma) upon rounding of the cells and gentle release from the flask. HUVECs were fixed in RPMI supplemented with 10 % FBS. Human Umbilical Derived Erythroid Progenitor line 2 cells^54^ (HUDEP-2; RIKEN) were maintained at 0.7-1.5×10^6^ cells ml^-1^ in HUDEP expansion media (SFEM, 50 ng/ml SCF, 3 IU/ml EPO, 10 µM DEX, 1% L-Glu, 1% Penstrep) and changed into fresh media containing 2x doxycycline (DOX) every two days. For differentiation we used a modified version of the CD34 differentiation protocol. 2-3×10^6^ cells were resuspended at 0.3-05×10^6^ cells ml^-1^ in HUDEP Phase I media (IMDM, 200 µg/ml Holotransferrin, 10 g/ml Insulin, 3 IU/ml Heparin, 3% Inactivated AB plasma, 2% FBS, 3 IU/ml EPO, 1 ng/ml IL-3, 10 ng/ml SCF, 1x Pen/Strep) with 1x DOX on day 0. On days 1 and 3 cells were counted, pelleted (5 min, 250 rcf, RT) and resuspended to 0.3-0.5×10^6^ cells ml^-1^ in fresh HUDEP Phase I media supplemented with 2x DOX. On day 5, cells were counted, pelleted and resuspended to 0.5×10^6^ cells ml^-1^ in HUDEP Phase II media (IMDM, 500 µg/ml Holotransferrin, 10 g/ml Insulin, 3 IU/ml Heparin, 3% Inactivated AB plasma, 2% FBS, 3 IU/ml EPO, 1x Pen/Strep) without DOX. On days 7 and 9 cells were counted, pelleted and resuspended to 0.5×10^6^ cells ml^-1^ in fresh HUDEP Phase II media. Cytospins and FACS was carried out as for CD34^+^ differentiation.

### HUDEP-2 genome editing

Prior to guide RNA (gRNA) design HUDEP-2 SNPs were genotyped by either Sanger or Next Generation sequencing at MRC WIMM Sequencing Facility with locus specific primers (Supplementary Table 3). For introduction of SNPs by CRISPR/Cas9 facilitated homology dependent repair (HDR), gRNAs were designed to cut in close proximity to the SNP of interest with the PAM overlapping the SNP where possible, additionally single stranded DNA donors (ssODN; IDT) were offset and thioated to promote integration and reduce degradation. To control for global effects on HUDEP-2 cells caused by CRISPR/Cas9 editing, gRNA and ssODN were designed for a homozygous SNP (rs4508712) with no GWAS associations (www.ebi.ac.uk/gwas), which was not within an erythroid regulatory element or expressed gene. HUDEP-2 genotype specific gRNAs (Merck) were cloned into pX458 plasmid backbone^68^ by the Genome Engineering Facility (WIMM, University of Oxford) and purified using Plasmid Midi Kit (Qiagen). pX458 (pSpCas9(BB)-2A-GFP) was a gift from Feng Zhang (Broad Institute) and is available from Addgene (plasmid #48138). HUDEP-2 cells were then transfected as previously described^55^. Briefly, ∼1×10^6^ cells were transfected with pairs of 5 µg gRNA plasmid and 4 µg ssODN (Supplementary Table 3) using Amaxa^TM^ Human CD34 Cell Nucleofector^TM^ Kit (Lonza) in the 2B-Nucleofector^TM^ on the U-08 setting. Cells were transferred to 2.5 ml HUDEP expansion media supplemented with 2x DOX and 7.5 µM RAD51-stimulatory compound 1 (RS-1, Sigma). After two days cells were pelleted (5 min, 250 rcf, room temp.) and resuspended in 2.5 ml HUDEP expansion media supplemented with 2x DOX with minimal light exposure. On day 3 cells were single cell sorted on a BD FACSAria Fusion flow cytometer (BD Bioscience) into terazaki plates containing 20 µl of expansion media (2x DOX). When colonies reached more than 30 cells they were transferred to a 96-well plate and expanded over two weeks with fresh media and DOX every 2 days until filling two to four wells of a 96-well plate. Half of the cells for each expanded clone were frozen (90% FBS, 10% DMSO) as a stock for recovery post genotyping. For genotyping we followed a 96-well barcoding approach with Next Generation sequencing^69^. Clonally amplified cells were first lysed (50 mM Tris, 1 mM EDTA, 0.5% Tween 20) and the targeted locus was amplified with primers containing a modified m13 adaptor sequence (Supplementary Table 3), the adaptor was then used to for priming with row and column specific primers in a second PCR to barcode each well. Finally, all wells from a single plate were pooled and prepared for sequencing with the NEBNext Ultra II DNA Library Prep kit for Illumina (New England Biolabs). Plates were multiplexed and sequenced on the MiSeq platform (Illumina) using 250 bp paired-end reads (Nano kit, v2 chemistry). Sequences were analysed using platescreen96^69^ (v4.0.4, <u>github.com/Hughes-Genome-Group/plateScreen96/releases</u>) to genotype clones. Screening was carried out to exclude for rs9349205 and rs4508712 clones which appeared homozygous due to microhomology driven large deletions (200-4,000 bp)^70, 71^, rather than through HDR editing, by PCR with locus specific primers (Supplementary Table 3). For rs10758656, two upstream heterozygous SNPs rs7870037 (+129 bp) & rs7855081 (+132 bp) allowed for the exclusion of loss of heterozygosity.

### Gene expression analyses

1-5×10^6^ cells were fixed in 1 ml TRI-reagent (Sigma), snap frozen and stored at −80°C for less than one year. RNA was extracted by addition of 0.1 ml 1-bromo-3-chloropropane, pipette mixing and separation in a Phase Lock gel Heavy tube (5Prime) and then precipitation with 1 µl of GlycoBlue and an equal volume (∼500 µl) isopropanol and centrifugation (10 min, 12,000 rcf, 4 °C). The RNA pellet was washed with 75% ethanol, resuspended in DEPC-treated water, and stored at −80°C for less than one year. For RT-qPCR RNA was treated with 2U of rDNase I (Invitrogen) and then 1 µg of RNA was used to generate cDNA using SuperScript III First Strand Synthesis SuperMix (Invitrogen) following the manufacturers’ instructions. Real-time RT-qPCR was performed on a StepOne Thermocycler (ThermoFisher) using Taqman Universal PCR Master Mix II (Life Tech) and commercially available expression assays (Supplementary Table 5; Life Tech). For RNA-seq total RNA was treated with Turbo DNase (Invitrogen) at 25°C for 60 min, then RNA was separated using phenol-chloroform isoamylalcohol and a PhaseLock Light-gel tube (5Prime). Treated RNA was precipitated at −80°C overnight with sodium acetate, glycoblue, and 75% ethanol, before centrifugation (12,000 rcf, 4°C), 75% ethanol wash and resuspension in DEPC-treated water. Globin and rRNA sequences were depleted from up to 5 µg of treated RNA using Globin-Zero Gold (Illumina), before PolyA selection with NEBNext Poly(A) mRNA Magnetic Isolation module (New England Biolabs), and indexing with NEBNext Ultra Directional RNA Library Prep Kit for Illumina (New England Biolabs) following the manufacturers’ instructions. RNA-seq libraries were quantified by qPCR (KAPA) prior to sequencing on the NextSeq platform (Illumina) with 39 bp paired-end reads. Reads were mapped to hg19 (Supplementary Table 6) using STAR^72^ (v2.4.2a; --outFilterMultimapNmax 1) and duplicates were filtered using samtools^73^ (v1.3; rmdup). For visualisation directional reads were normalised to RPKM using deepTools^74^ with no windowing (v2.2.2; bamCoverage --binSize 1 --normalizeUsingRPKM). Uniquely mapped reads were analysed in DESeq2^61^ using variance stabilising transformation and exclusion of genes lacking 5 total reads. Expressed genes were classed as having more than log2(FPKM) greater than −5.

### Chromatin conformation capture and target gene identification

For chromatin conformation, 1-2×10^7^, H1-hESC, HUVEC or erythroid cells were crosslinked with 2% formaldehyde which provides optimal *cis/trans* ratios and digestion efficiencies^58^. For each cell type triplicate 3C libraries were prepared using *Dpn*II and standard methods^23^ with the following modifications: no douncing was performed, all spins were performed at 300 rcf, and after ligation intact nuclei were pelleted (15 min, 300 rcf), supernatant was discarded, and nuclei were resuspended in 300 µl Tris-EDTA (TE; Sigma) for phenol chloroform extraction. Digestion efficiency was determined by RT-qPCR with TaqMan and custom oligonucleotides (Supplementary Table 5), and ligation efficiency qualitatively determined by gel electrophoresis. Only 3C libraries with >70% digestion efficiencies were used. 3C libraries were sonicated to 200 bp in a Covaris S220 and indexed with NEB Next Illumina library Prep reagents (NEB). Enrichment for specific viewpoints was performed with 70mer biotinylated oligonucleotides designed using CapSequm^75^ (http://apps.molbiol.ox.ac.uk/CaptureC/cgi-bin/CapSequm.cgi). Double capture was performed in multiplexed reactions with pools of oligonucleotides targeting either promoter proximal (within 5 kb of a transcription start site) or promoter distal (enhancer centric) *DpnII* fragments (Supplementary Table 7) following the described method^23^ with each oligonucleotide at a working concentration of 2.9 nM. Captured 3C libraries were sequenced on the NextSeq platform (Illumina) with 150 bp paired-end reads. Reads were mapped and analysed using CCseqBasic5 (github.com/Hughes-Genome-Group/CCseqBasic5) as previously described^76^ with the following custom settings (--bowtie2 --globin 2). Briefly, CCseqBasic5 trims adaptor sequences, flashes read pairs, *in silco* digests fragments and uses bowtie2 to map reads before identifying capture and reporter reads. After primary analysis replicates were compared using the comprehensive CaptureCompare software (github.com/Hughes-Genome-Group/CaptureCompare). CaptureCompare normalises *cis* reporter counts per 100,000 *cis* reporters, generates per fragment mean counts for each cell type, calculates difference in mean interactions between cell types, compares differences in raw interaction counts per fragment using DESeq2^45^ as previously described^23, 77, 78^, and provides input for peaky interaction calling^47^. Interaction calling using peaky was run with default settings (omega −3.8) and interactions were filtered based upon the Marginal Posterior Probability of Contact (MPPC) within local interaction domains (MPPC > 0.01) or within 1 Mb of the viewpoint (MPPC > 0.1) and assigned to either Refseq transcription start sites (tss) or variants within 500 bp of the significantly interacting fragment. Target genes were first identified as those having a tss within 5kb of an intersecting variant (promoter proximity), being within 500 bp of a significantly enriched erythroid fragment (FDR <0.05) or within 500 bp of a peaky identified interaction. Candidate genes were subsequently filtered for detectable erythroid expression (log2(FPKM>-5)) using RNA-seq. 24 benchmarking genes most likely to be effectors were identified based on published functional data (*IKZF1, KIT, TAL1, RBM38, SMIM1, CD164, CCND3, MYB, HBA1, HBA2, BCL11A, JAK2*)^5, 12–14, 16, 42, 79^, presence in the Oxford Red Cell Panel for rare inherited anaemia (*KLF1, TFRC, ANK1, HK1, SCL4A1*)^80^, containing mutations causing hemochromatosis (*TFR2*)^81^, having an erythroid eQTL (*ATP2B4*)^15^, and causing altered RBC phenotypes in mouse and zebrafish (*FBXO7, CCNA2, miR-181a, PIEZO1, AKAP10, CITED2*)^82–88^.

### Chromatin IP, ATAC-seq, and data processing

For ChIP-seq, chromatin was crosslinked with 1% formaldehyde (Sigma) by the addition of 1 ml 10x crosslinking buffer (50 mM HEPES, 1 mM EDTA, 0.5 mM EGTA, 100 mM NaCl, 10% formaldehyde) to 10^7^ cells in 9 ml of media and incubation at room temperature for 10 minutes. Crosslinking was quenched with 130 mM glycine, and cells were washed with cold PBS before snap freezing pelleted cells. Fixed material was stored at −80°C for less than 12 months. Chromatin immunoprecipitation was performed using Agarose ChIP Assay Kit (Merck Millipore). Briefly,10^7^ cells were lysed by incubation on ice with 130 µl lysis buffer for 15 minutes. Lysed cells were transferred to Covaris microtubes and sonicated on the Covaris S220 (Duty cycle: 2%, Intensity: 3, Cycles per burst: 200, Power mode: Frequency sweeping, Duration: 480 sec, Temp.: 6°C) to generate 200-400 bp fragments. Insoluble material was removed by centrifugation (15,000 rcf, 15 min, 4°C) and soluble material was diluted to 4 ml with dilution buffer. Immunoprecipitation was performed by incubation of 2 ml diluted chromatin (equivalent to 5×10^6^ input cells) with antibodies for H3K4me1 (3 µg ab195391, lot: GR304893-2; AbCam), H3K4me3 (1 µl 07-473, lot: 2664283; Millipore), H3K27ac (0.3 µg ab4729, lot: GR3205523-1; AbCam), CTCF (10 µl 07-729, lot: 2836929; Millipore) or GATA1 (∼7.2 µg ab11852, lots: GR208255-9, GR290167-7, GR239235-8; AbCam) overnight. Chromatin binding to Protein A/agarose slurry, washes and elution were performed according to the manufacturer’s instructions. DNA was purified by phenol-chloroform extraction with PhaseLock tubes (5Prime) and ethanol precipitation with NaOAc, and 2 µl GlycoBlue (Invitrogen). ChIP enrichment was determined by RT-qPCR (Supplementary Table 5) prior to addition of sequencing adaptors using NEBNext Ultra II DNA Library Prep kit for Illumina (New England Biolabs). ATAC-seq was performed as previously described^76, 89^ using two technical replicates with 7×10^5^ cells. ChIP-seq and ATAC-seq libraries were quantified by RT-qPCR with the KAPA Library Quantification Complete Kit (KAPA) prior to sequencing on the NextSeq platform (Illumina) with 39 bp paired-end reads. ATAC-seq, DNaseI-seq and ChIP-seq reads were mapped to the hg19 genome using NGseqBasic^90^ (V20; --nextera --blacklistFilter --noWindow) which utilises bowtie. Sequence depth and mapped reads for each sample are provided (Supplementary Table 6). Published GEO repositories^24, 66, 91–98^ were downloaded for ATAC-seq and DNaseI-seq from HSC, CMP, MEP, MPP, Ery (GSE75384), and HUVEC (GSM736575, GSM736533), and ChIP-seq for SCL/TAL (GSE95875, GSE93372, GSE42390, GSE70660, GSE59087, GSE52924), GATA1 (GSE32491, GSE36985, GSE107726, GSE29196), NF-E2 (GSE95875), BACH1 and MAFK (GSE31477), and SPI1/PU1 (GSE70660), and were analysed by the same method. For visualisation, PCR-duplicate filtered replicates were merged using samtools^73^ (v1.3) and converted to bigwigs with minimal smoothing using deepTools^74^ (v2.2.2; bamCoverage -- binSize 10 --normalizeUsingRPKM --minMappingQuality 30).

### Imputation and in silico analysis of variants

The original 75 anaemia index SNPs were imputed with HapMap Phase 2 which is lower resolution than the 1000 Genomes Project Phase 3 dataset^11, 99^. Therefore variants in linkage disequilibrium (LD) with index SNPs were identified using the rAggr proxy search online tool (<u>raggr.usc.edu</u>) with default settings (r^2^≥0.8, distance limit: 500 kb, population panels: All European, All South Asian) for the 1000 Genomes Project Phase 3 database^100^, which generated 6,420 variants. LD variants for Astle *et al.* (2016) were provided by Lisa Schmunk, Tao Jiang, and Nicole Soranzo (University of Cambridge). Summary statistics for Malaria^101^, Multiple Sclerosis^25^, Inflammatory Bowel Disease^27^, Type 1 Diabetes^26^, Type 2 Diabetes^102^, Intelligence^28^, and Central Corneal Thickness^29^ were downloaded from the NHGRI-EBI GWAS Catalog^1^ (www.ebi.ac.uk/gwas). For comparison of linkage between index SNPs from van der Harst *et al.* (2012)^11^ and Astle *et al*. (2016)^10^ we used LDmatrix on the LDlink web tool^103^ (http://ldlink.nci.nih.gov/) for European populations as this matched the cohort of the later study. Variants were intersected with peak calls from ATAC-seq or DNaseI-seq for each cell type of interest using bedtools^104^. Enrichment was calculated as the -log(p-value) of a binomial cumulative distribution function b(*x*; *n*, *p*), describing the probability of *x* or more successes from *n* Bernoulli trials, with the probability of success for each trial being *p.* P-values were calculated using the R function pbinom (lower.tail=FALSE) where *x* was the number of intersecting variants, *n* was the total number of variants and *p* was the total number of base-pairs within cell specific peaks divided by the hg19 uniquely mappable base-pairs (2,644,741,479 bp). Variants within the exons and introns of expressed coding genes were tested for predicted damaging effects on coding ANNOVAR^17^ or splicing SPIDEX^18^ (z-score ≥1.65) and SpliceAI^19^ (AI score ≥0.2). Variants within open chromatin were assessed for potential damage to transcription factor binding footprints using Sasquatch^20^ (7-mer, WIMM Fibach Erythroid, Exhaustive). Variants within open chromatin were further classified based on their predicted effect on chromatin accessibility using a deep convolutional neural net: deepHaem^21^. Model architecture and data encoding were adapted from DeepSEA^36^ with the following modifications: the number of convolutional layers was increased from three to five; batch normalisation was excluded as it did not improve convergence. The network was re-implemented in python using tensorflow (v1.8.0; https://www.tensorflow.org/about/bib). The ENCODE data compendium previously used^36^ was supplemented with ATAC-seq and CTCF ChIP-seq data from erythroid differentiations generated for this work, DNAseI-seq^20^, and ATAC-seq from sorted progenitor populations^24^.

Full model details and architecture are available on GitHub (https://github.com/rschwess/deepHaem).

### Chromatin segmentation and enhancer based PCA analysis

To ensure identification of all ATAC-seq peaks a combination of the traditional MACS2 approach^105^ (v2.0/10 callpeak -B -q 0.01) and digital signal processing with Ritornello^106^ (v2.0 default settings) was used. Peak summits from both calls were extended to 500 bp and intersected with bedtools^104^ (v2.25.0), and filtered for high ploidy regions in MIG viewer^107^ to form peak calls for each cell type (Supplementary Tables 8a-p). Chromatin segmentation was performed using the GenoSTAN^31^ hidden Markov model (HMM) which allows a more fine-tuned analysis than ChromHMM^108^ as it uses continuous rather than binary signal counts. Segmentation used a peak centric approach, rather than signal across the whole genome, with triplicate H3K4me1, H3K4me3, H3K27ac, and CTCF from day 10 of *ex vivo* CD34 differentiation. Read coverage of each mark was calculated (deepTools v2.4.2) for 1 kb windows over open chromatin peaks (bedtools merge -d 10) to capture histone modifications. The HMM model was trained using Poisson log-normal distributions with 20 initial states. These were manually curated to eight final states based on similarity of chromatin signature. For Principle Component Analysis (PCA) trajectory plotting combined peak calls from sorted hematopoietic populations covering 176,135 open chromatin regions not within 2 kb of transcription start sites were first used to generate a PCA map of erythroid differentiation from sorted populations of HSC, MPP, CMP, MEP and Erythroid populations^24^. Reads within peaks were normalised (R scale) and the PCA was calculated using the R function prcomp. The read counts from *ex vivo* differentiated cells within the same peak set were then used to calculate sample mapping onto PC1 and PC2, and thus to map differentiation timepoints onto the differentiation trajectory. Heatmaps of intersected peaks were generated with pheatmap^109^ (v1.0.8) using z-normalised counts of reads per basepair from all identified peaks. For enhancer activity, peak calls were extended by 250 bp in both directions (bedtools slop) to account for the spreading nature of H3K27ac ChIP-seq signal, enhancers were then ranked based on reads per base pair. To determine the point of inflection between low and high acting enhancers H3K27ac read counts were transformed so that the highest value equalled the number of ranked peaks. The point of inflection where the gradient of the curve became greater than one was used to define low and high enhancer activity. The gradient was calculated for each peak as a window based on the local linear gradient of ±200 peaks.

## DECLARATIONS

### Ethics approval and consent to participate

All donors provided consent to participate in this study. Blood was collected with ethics approval by the NHS Health Research Authority (National Research Ethics Service, REC 03/08/097) and stored according to Human Tissue Authority guidelines (License 12433).

### Consent for publication

Not applicable

### Availability of data and materials

Sequence reads and processed data generated for this work have been archived with GEO; H1-hESC ATAC-seq (GSE124853), Immune cell ATAC-seq (GSE125164), HUDEP-2 data (GSE125715), primary human erythroid data (GSE125926) and processed data are available via a UCSC hub (http://sara.molbiol.ox.ac.uk/public/hugheslab/GWAS_Pipeline/hub.txt).

### Author Contributions

D.J.D., J.K., J.O.J.D., P.H., G.L., J.A.T., S.T., V.J.B., D.R.H., and J.R.H. designed and planned experiments. D.J.D., S.J.H., L.N., C.S., M.E.G., P.P.H., M.C.S., J.L.D., A.J.C., C.A.D., M.S.C.L., A.M.O., Y.B., A.J.K., P.H., and C.F. performed experiments. D.J.D., R.S., J.M.T., C.Q.E., S.J.M., J.R.J.I., and R.C.B. processed and analysed data. R.K., and Y.N. provided essential reagents. D.J.D., J.A.T., D.R.H, J.R.H. wrote the manuscript.

## Acknowledgements

We appreciate the time and support of Kevin Clarke, Sally-Anne Clarke and Paul Sopp (WIMM Flow Cytometry Facility, Oxford University) during cell sorting.

---

**Supplementary Information for:** An integrated platform to systematically find causal variants and genes for polygenic human traits.

**Supplementary Fig. 1.**
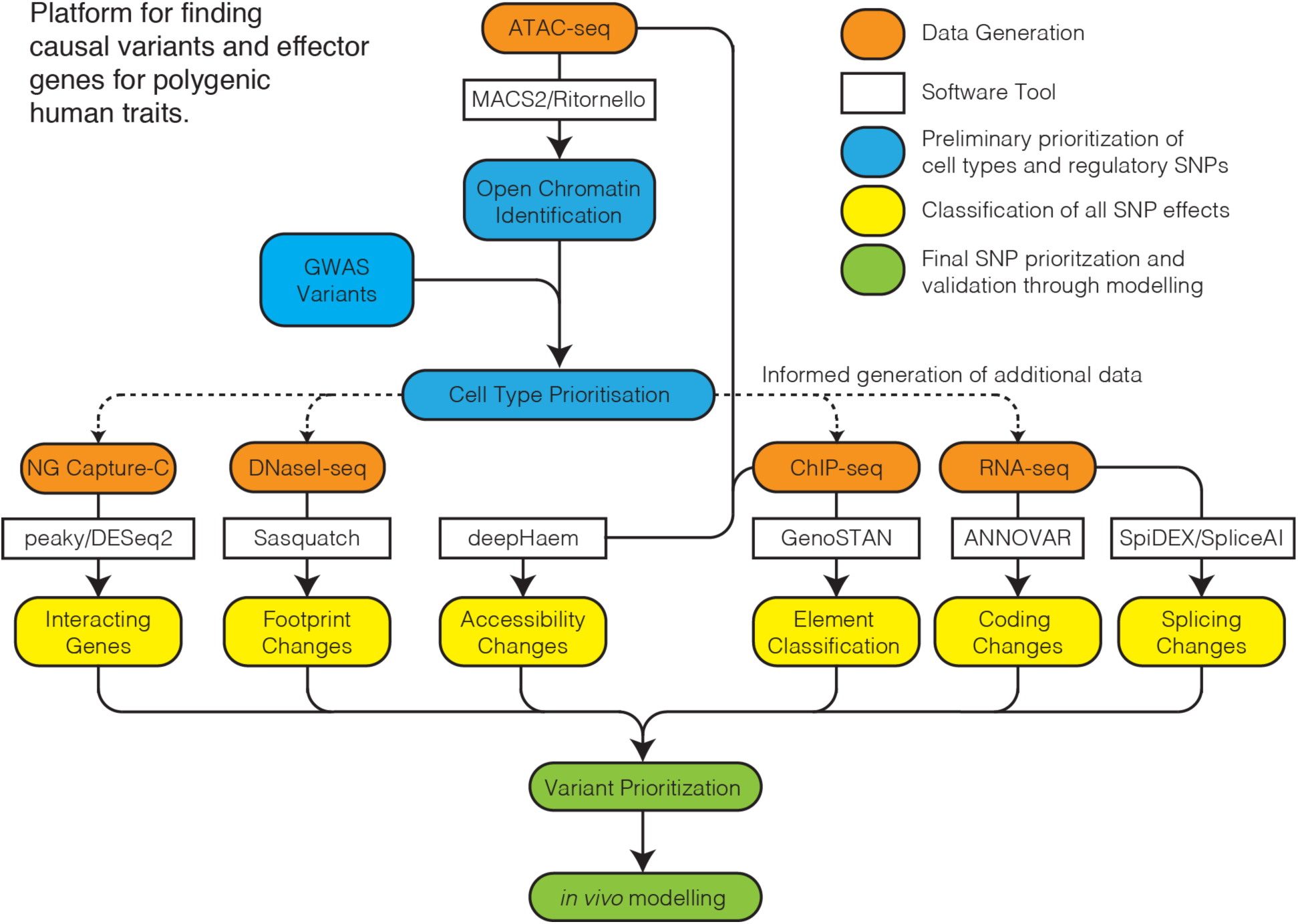
Flowchart of an integrated platform for dissecting multigenic traits using a multi-omics and machine learning approach. ATAC-seq is generated for multiple candidate cell-types and open chromatin regions are identified by MACS2 and Ritornello peak calling. Open chromatin regions for all cell types are intersected with GWAS variants to identify the key cell type(s) for further study. Additional -omics data is then generated in the key cell type(s) to characterise open chromatin regions (ChIP-seq), transcription factor footprints (DNaseI-seq), chromatin interactions (NG Capture-C) and expressed genes (RNA-seq). Note, Omics data is integrated through a suite of traditional and machine learning software (GenoSTAN, Sasquatch, deepHaem, peaky, DESeq2, ANNOVAR, SpiDEX, SpliceAI) to determine candidate causative variants and effector genes. Sasquatch currently has 314 available datasets for analysis, so DNaseI-seq would only need to be generated if the relevant cell type was not available. This wholistic approach allows the prioritization of variants for which can then be tested functionally by *in vivo* modelling.

**Supplementary Fig. 2.**
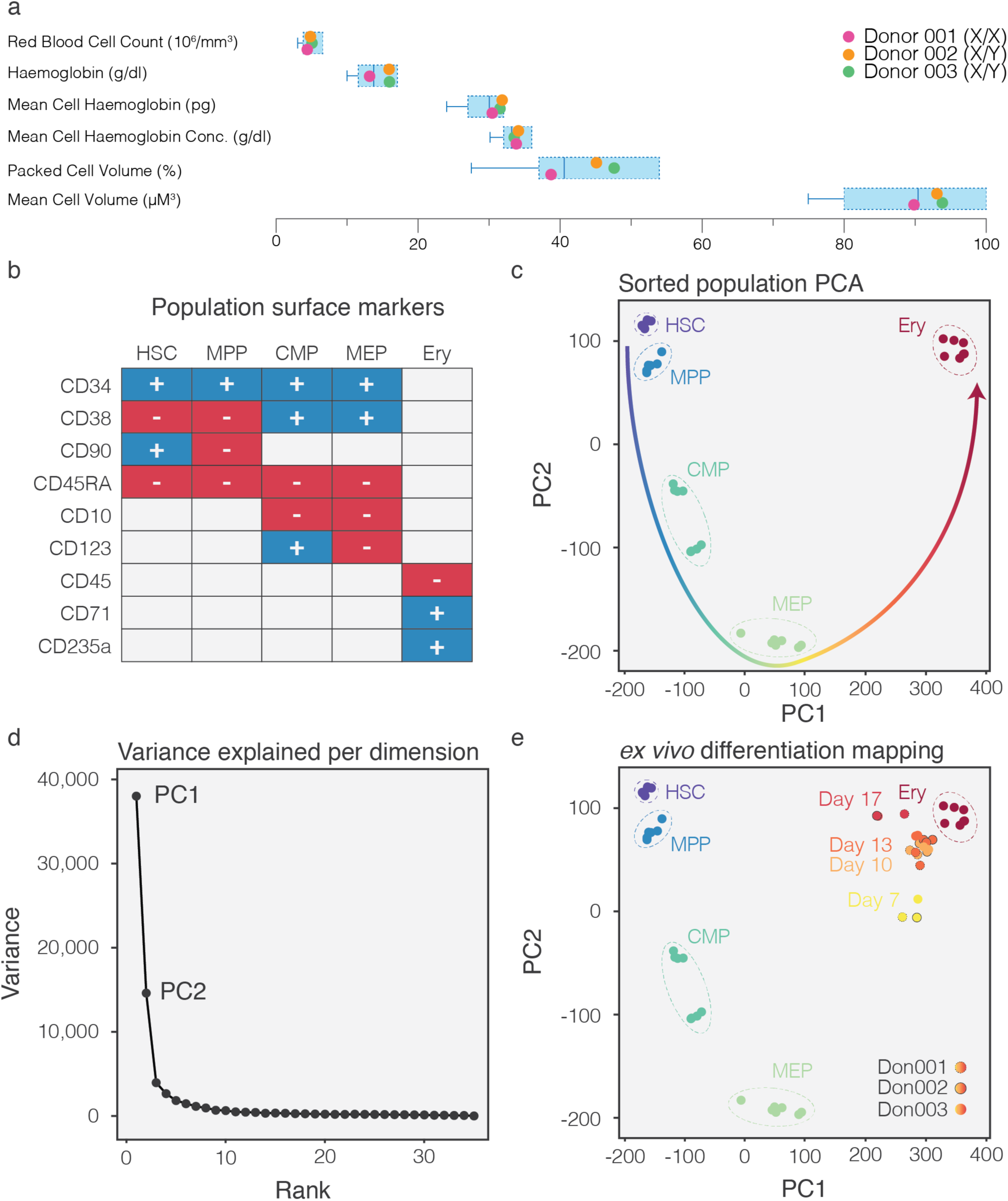
Differentiation of CD34^+^ cells from healthy donors. **a,** Whole cell counts for three healthy donors were compared against Oxford University Hospital, NHS Foundation Trust Normal ranges for adults (boxes) and GWAS exclusion values (whiskers) prior to differentiation. **b,** Cell surface markers used to define populations from published sorted cell population ATAC-seq (Corces *et al.* 2016). **c,** Principal Component Analysis (PCA) of ATAC-seq peaks outside 2 kb of transcription start sites maps the erythropoietic differentiation trajectory using the first two principal components (PC) which account for the majority of the variance (**d**). **e,** ATAC-seq from *ex vivo* CD34+ differentiation can be used to map progress through erythropoiesis by calculating PC1 and PC2 and shows high reproducibility between donors and replicate differentiations.

**Supplementary Fig. 3.**
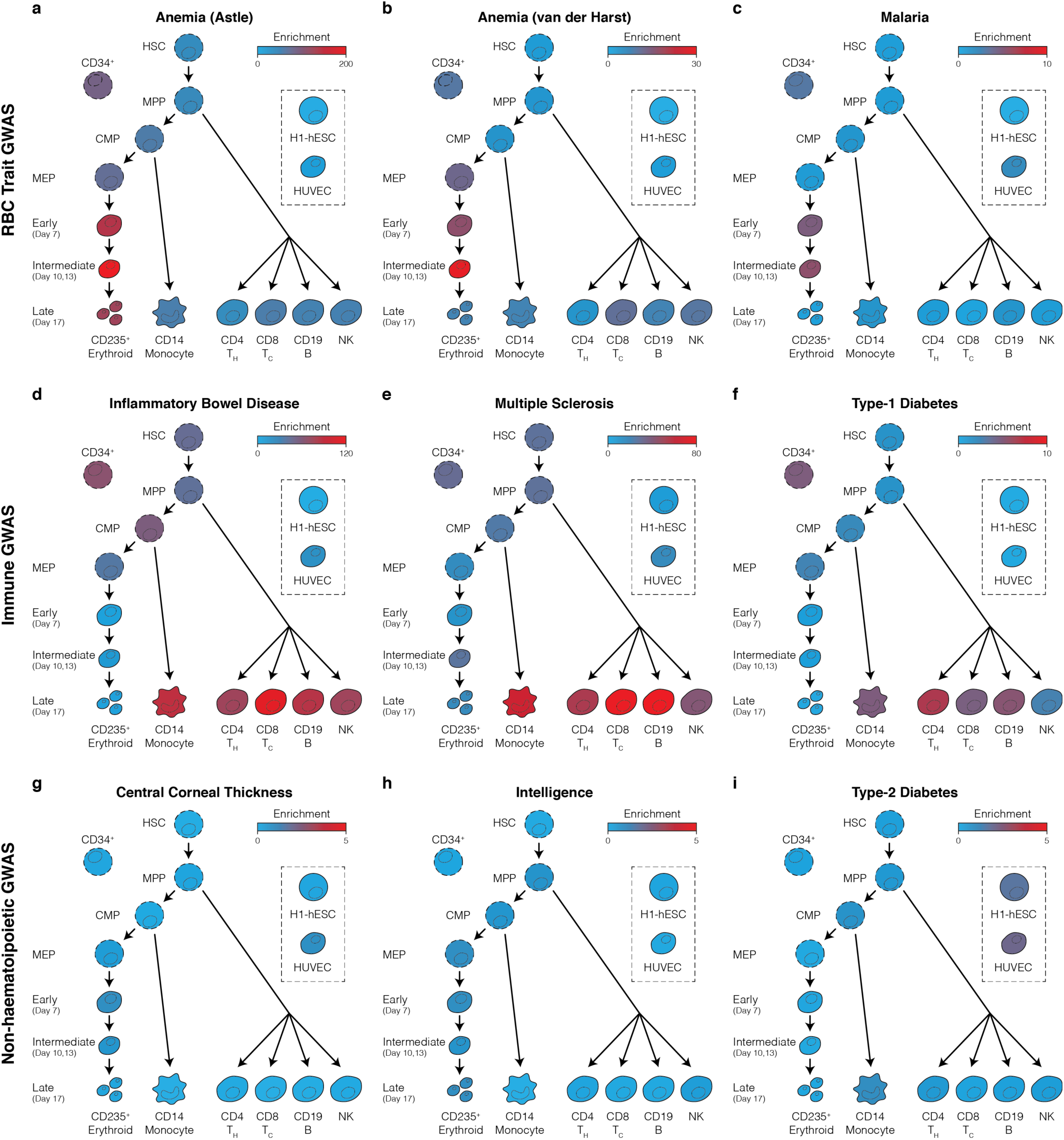
ATAC-seq peak intersections show disease specific profiles. Schematic of human haematopoiesis with enrichment (-log(p) of a cumulative Binomial Distribution) for red blood cell (**a-c**), monocytes and lymphocytes (**d-f**) and non-haematological (**g-i**) GWAS associated variants within open chromatin of haematopoietic stem cells (HSC), multi-potent progenitors (MPP), common myeloid progenitors (CMP), megakaryocyte-erythroid progenitors (MEP), erythroid cells from *in vitro* culture, CD14 monocytes, CD4 helper and CD8 cytotoxic T-cells, CD19+ B-cells, natural killer cells (NK), human embryonic stem cells (H1-hESC) and human umbilical vein endothelial cells (HUVEC).

**Supplementary Fig. 4.**
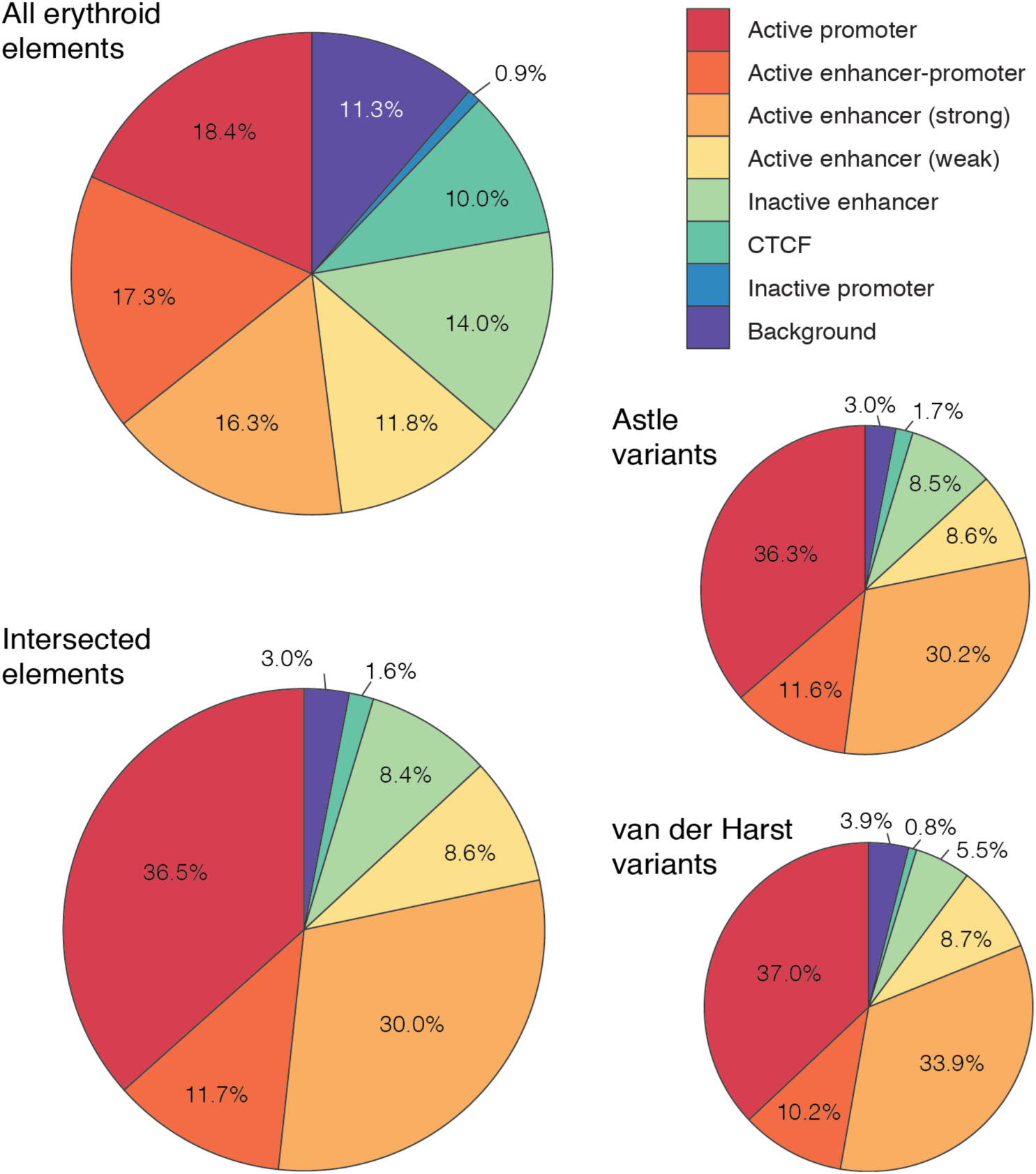
RBC Trait containing ATAC-seq peaks are enriched for strong enhancers and promoters. GenoSTAN uses a hidden Markov model and was implemented to classify defined chromatin regions into “states” using ChIP-seq read counts. Erythroid open chromatin regions were subclassified into 7 functional classes and a background class using H3K4me3 (promoter mark), H3K4me1 (enhancer mark), H3K27ac (active transcription mark) and CTCF (boundary element) ChIP-seq. Following intersection of open chromatin regions from two RBC trait GWAS studies, the re-distribution of GenoSTAN states of open chromatin regions containing variants was determined. Intersected elements are enriched for the Promoter and Strong Enhancer (high levels of H3K27ac) classes.

**Supplementary Fig. 5.**
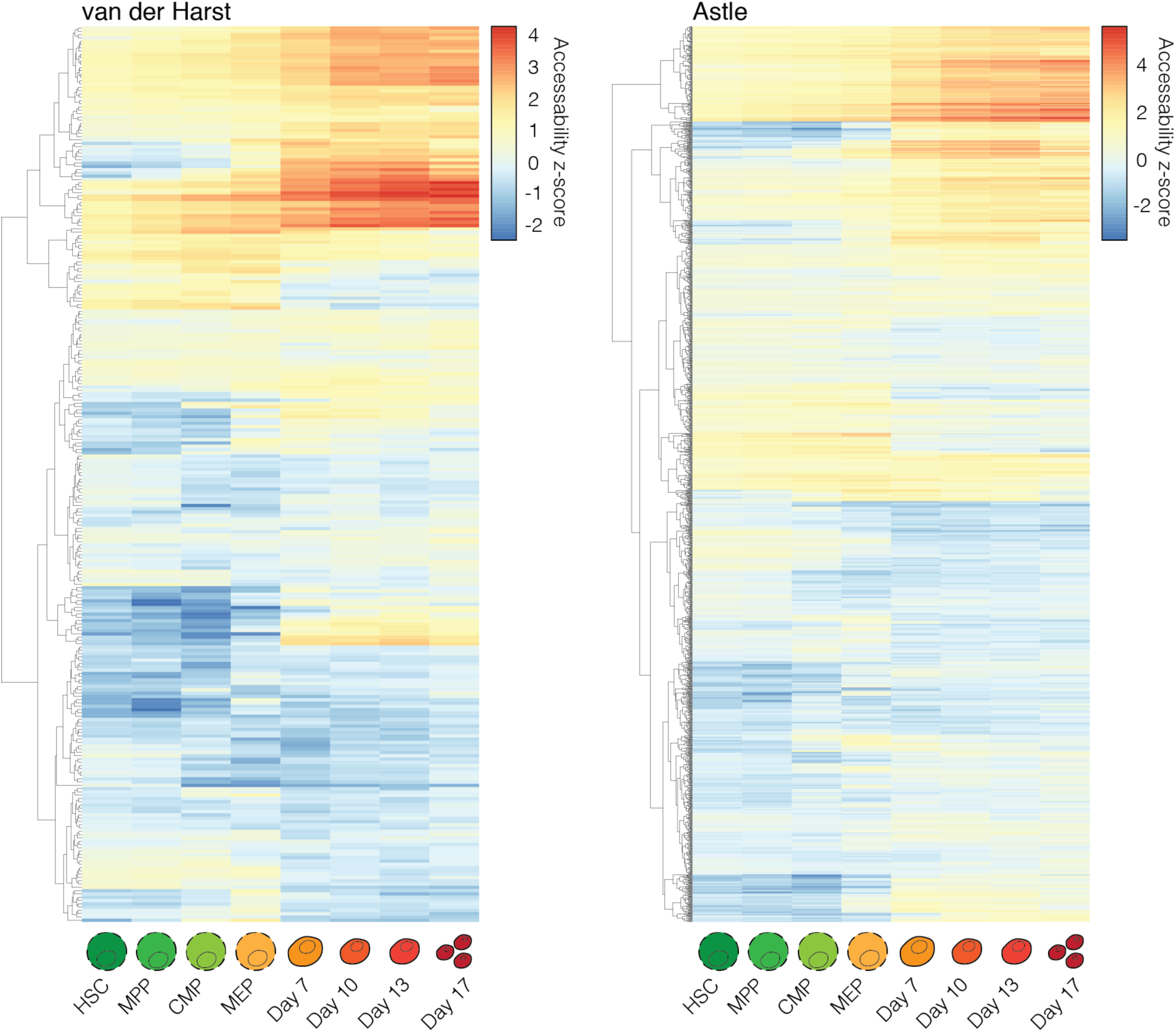
Intersected peaks show multiple trajectories of openness. pHeatmap clustering of z-score normalised ATAC-seq read counts for RBC variant containing peaks from haematopoietic stem cells (HSC), multi-potent progenitors (MPP), common myeloid progenitors (CMP), megakaryocyte-erythroid progenitors (MEP), *in vitro* differentiated erythroid cells (Day7-17).

**Supplementary Fig. 6.**
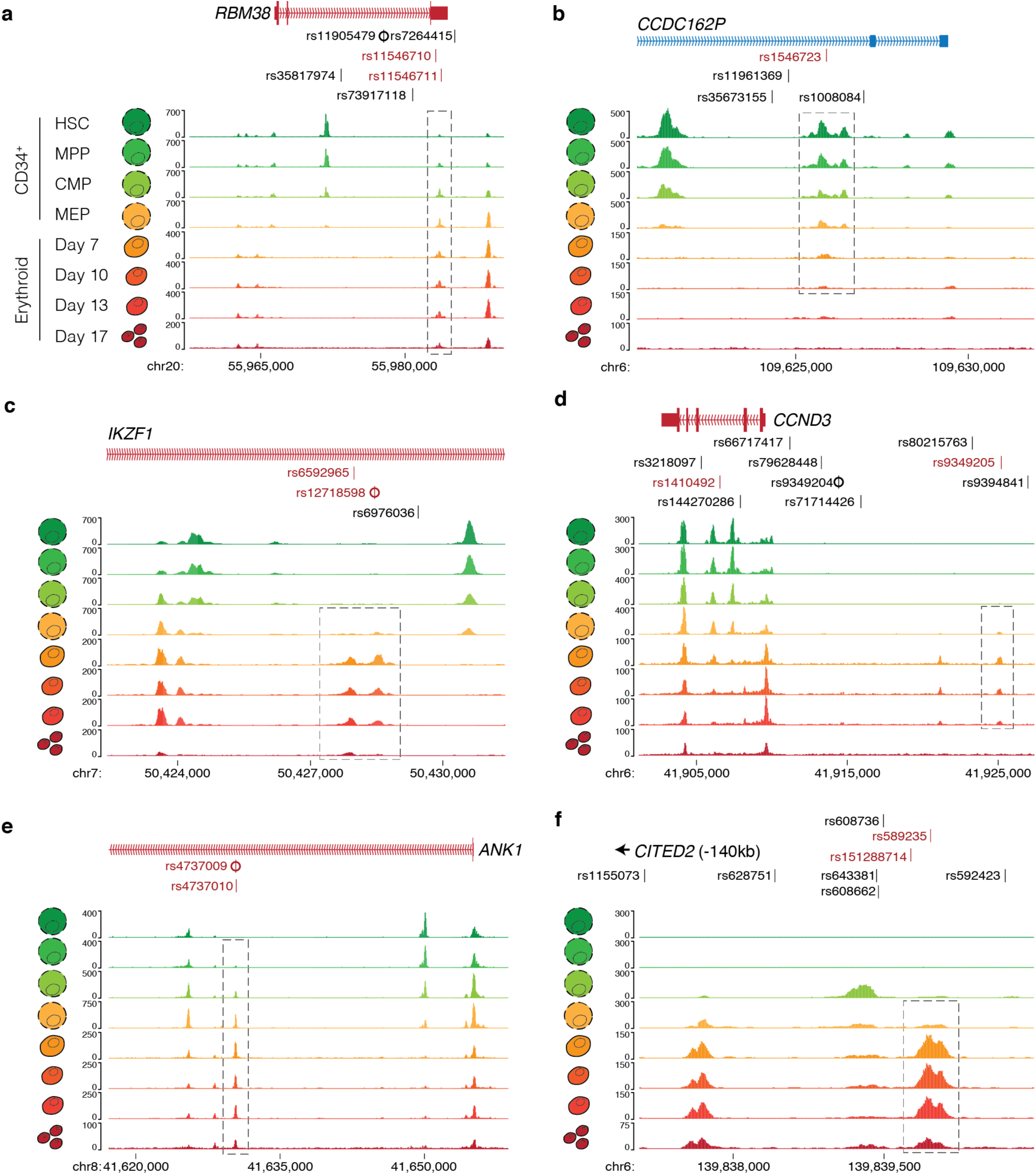
Examples of ATAC-seq peak intersection trajectories. ATAC-seq from haematopoietic stem cells (HSC), multi-potent progenitors (MPP), common myeloid progenitors (CMP), megakaryocyte-erythroid progenitors (MEP), *in vitro* differentiated erythroid cells was intersected with trait associated variants. Trajectories of intersection included continual (**a**), progenitor (**b**), transient (**c,d**) and terminal (**e,f**) trajectories (boxed). Variants within open chromatin are red, the index SNP is marked with a circle.

**Supplementary Fig. 7.**
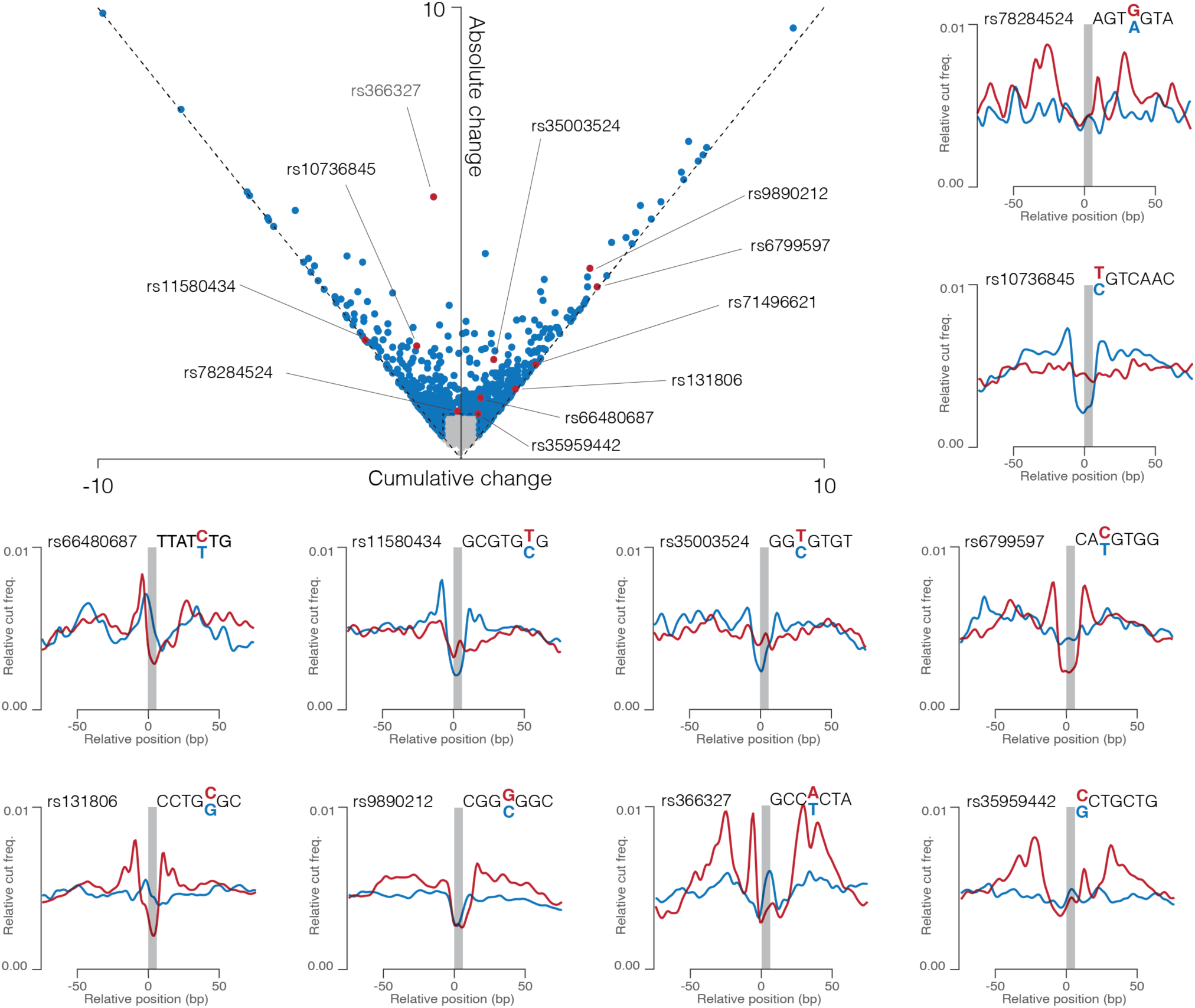
Sasquatch predicts changes in transcription factor footprints. Sasquatch generates meta-genomic footprints of 7-mer motifs using DNaseI signal from open chromatin regions to measure relative cut frequency. For variants in open chromatin, footprints and relative cut frequencies were generated with all seven possible 7-mer sequences for both alleles. Cumulative change and Absolute change for a variant are calculated by comparing the difference in relative cut frequency for each 7-mer and calculating the cumulative (directional) and absolute (total) change. Positive cumulative change scores can be interpreted as loss of a footprint, while negative values represent a gain of a footprint. Absolute change scores much larger than the cumulative change score (e.g. rs366327) represent a change in footprint type rather than a simple loss/gain. Variants with a cumulative change ≥0.5 or an absolute change ≥1.0 were considered damaging (blue dots). Representative footprints are shown for a range of SNPs (red dots).

**Supplementary Fig. 8.**
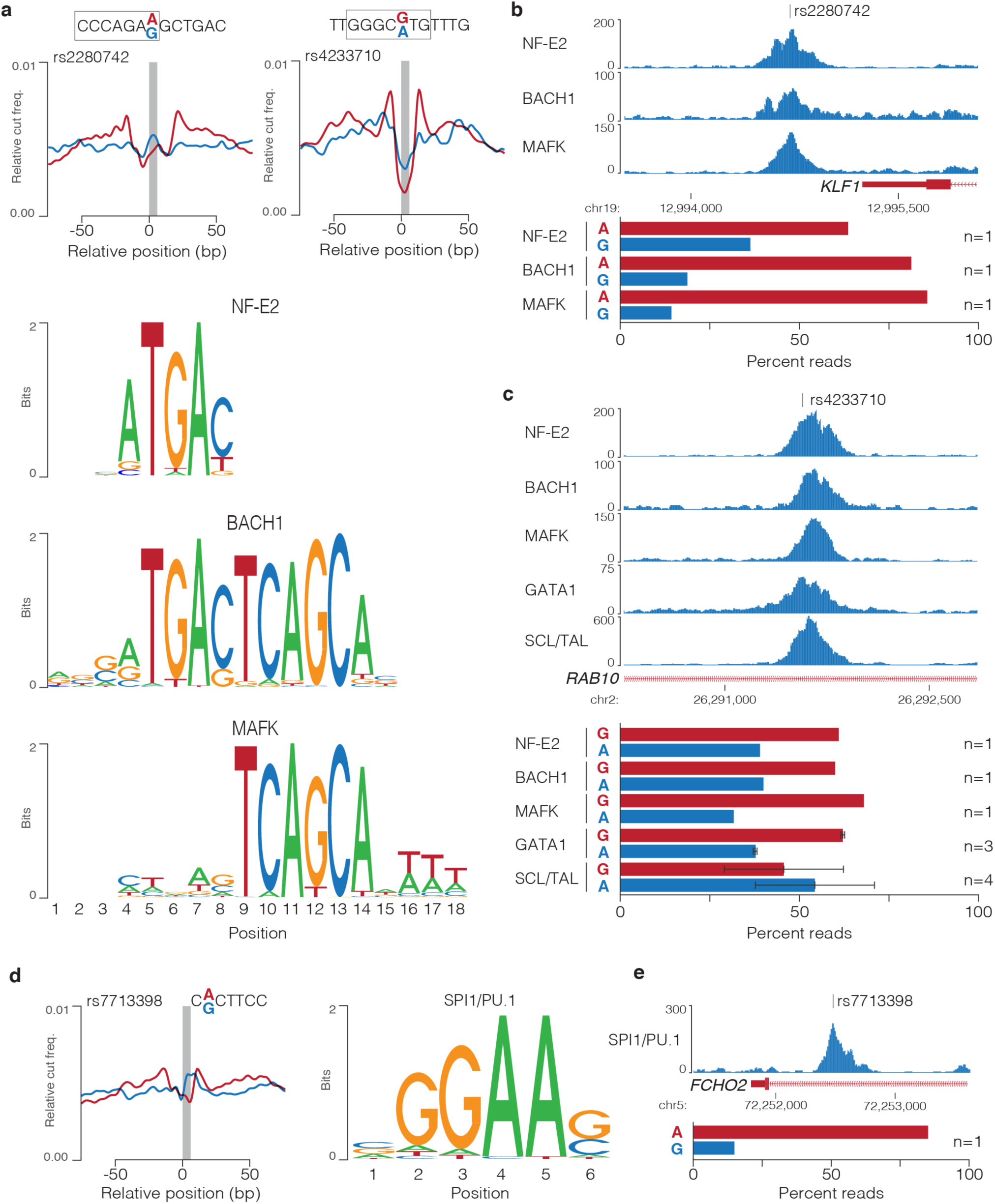
) Sasquatch predicted changes correlate with skew in binding of multiple transcription factors. **a,** rs2280742 and rs4233701 lie within 7-mer motifs (boxed) that are adjacent to binding motifs for NF-E2, BACH1 and MAFK and are predicted to alter DNaseI footprints at these sites. ChIP-seq for NF-E2, BACH1, MAFK, GATA1, and SCL/TAL over rs2280742 (**b**) and rs4233701 (**c**) show allelic skew in multiple factors as shown by percent of reads containing either allele. Error bars depict standard error of the mean. **d,** rs7713398 alters the predicted DNaseI footprint for a 7-mer motif which contains an SPI1/PU.1 consensus motif. **e,** ChIP-seq of SPI1/PU.1 shows allelic skew over rs7713398, as shown by percent of reads containing either allele.

**Supplementary Fig. 9.**
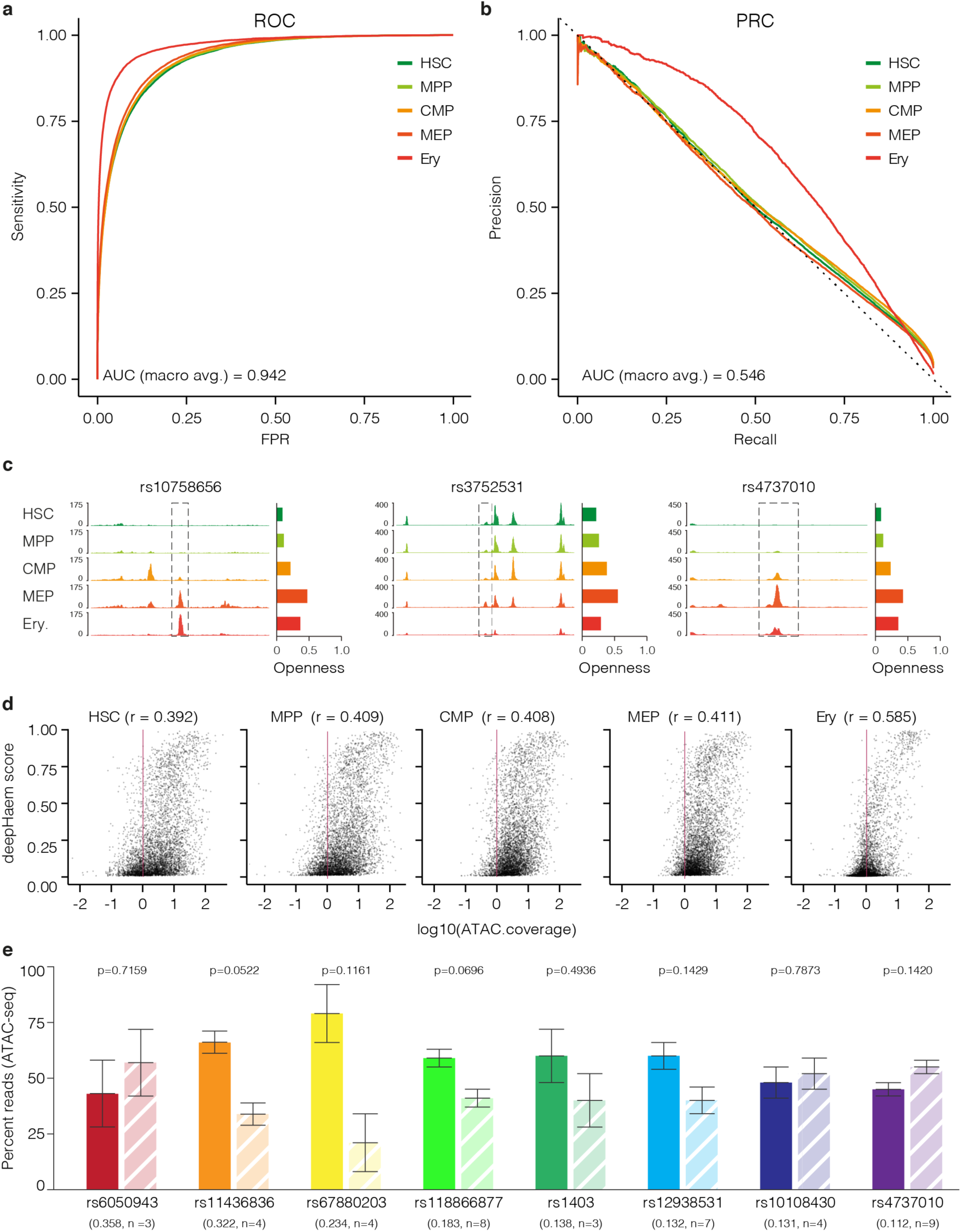
deepHaem prediction chromatin openness throughout erythropoiesis. Receiver operating characteristic (ROC) curve (**a**) and Precision-Recall Curve (PRC) (**b**) for deepHaem in haematopoietic stem cells (HSC), multi-potent progenitors (MPP), common myeloid progenitors (CMP), megakaryocyte-erythroid progenitors (MEP) and erythroid (Ery). **c**, FPKM normalised ATAC-seq around three SNPs predicted to be damaging by deepHaem, with openness scores shown for the hg19 reference sequence calculated for the 1kb box (dashed lines) centred on the SNP. **d**, Pearson correlation between deepHaem open chromatin scores and ATAC-seq coverage in corresponding cell types in a random sampling (1/50^th^) of MACS2 identified peaks. **e,** Mean percentage of day 10 and day 13 erythroid ATAC-seq reads on either the reference (dark bars) or variant (light dashed bars) allele from heterozygous individuals with a minimum of 5 reads. Error bars depict the standard error of the mean. deepHaem change score and number of independent replicates (including multiple donors and differentiations) are shown in parenthesis. p-values shown are for a ratio paired t-test.

**Supplementary Fig. 10.**
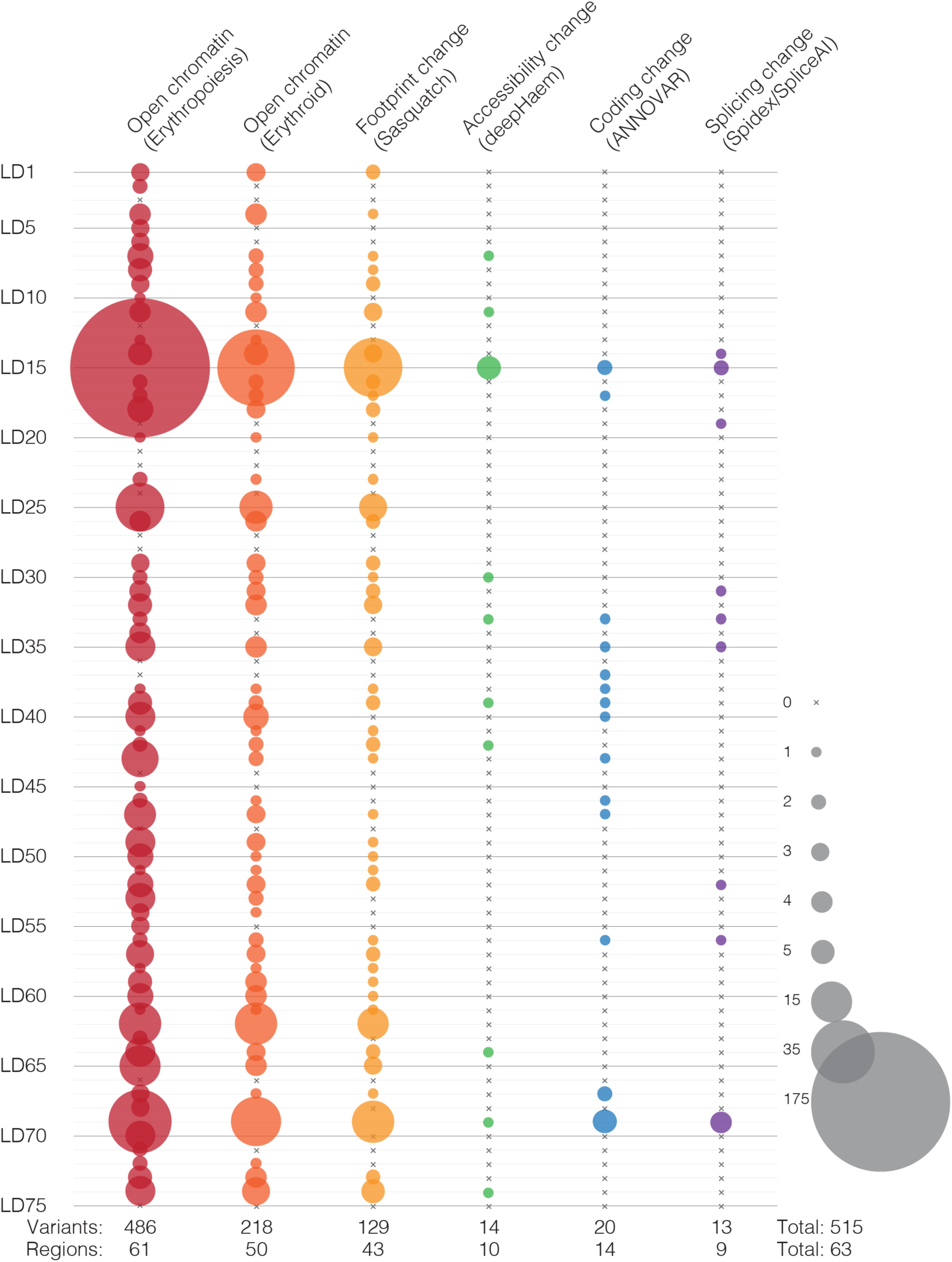
Pipeline summary for prioritization GWAS variants. RBC GWAS variants in linkage disequilibrium (LD) at 75 chromosome regions (n=6,420) were examined for presence in open chromatin, and putative changes in DNaseI footprint, chromatin accessibility, coding sequence, splicing potential. The total number of variants in each class is shown at the bottom. Circle size indicated the number of variants included for each class in a chromosome region.

**Supplementary Fig. 11.**
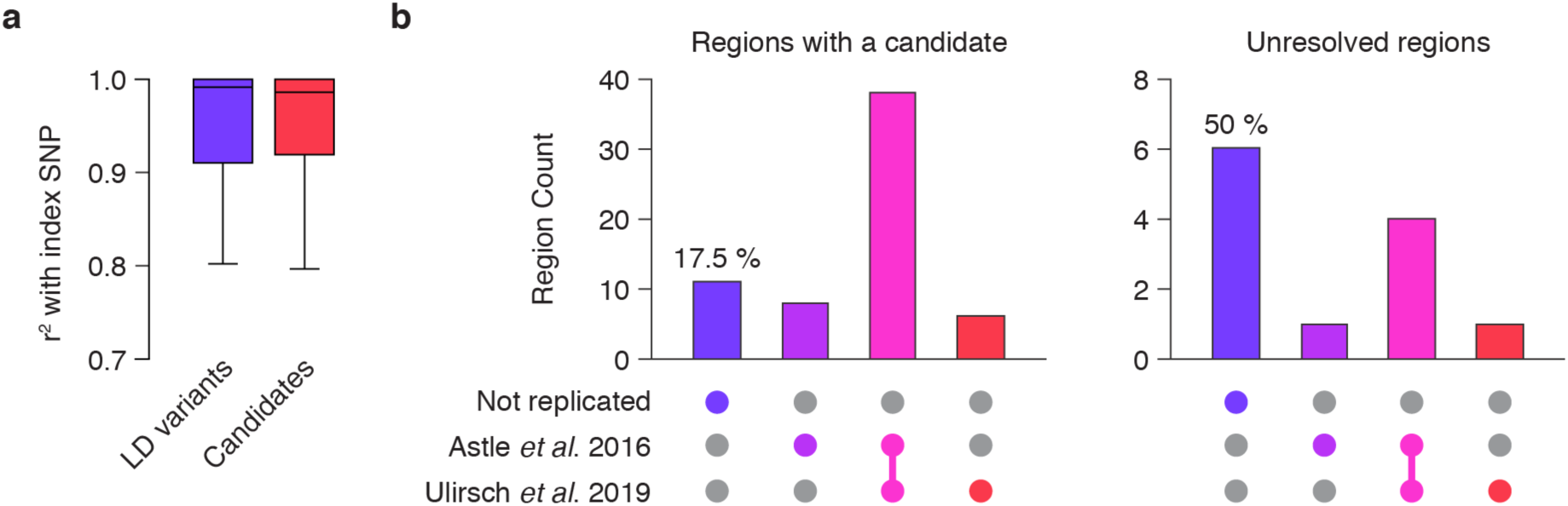
Analysis of resolved and unresolved regions. **a,** r^2^ for linkage disequilibrium analysis of RBC trait index SNPs with all analysed variants (n=6,420) and candidate variants identified by the platform (n=515). **b,** Counts of regions where index SNPs identified in van der Harst *et al.* (2012) was replicated in either of two subsequent RBC trait GWAS. Regions are separated into those with candidate variants following analysis in the described platform (n=63), and those without (n=12).

**Supplementary Fig. 12.**
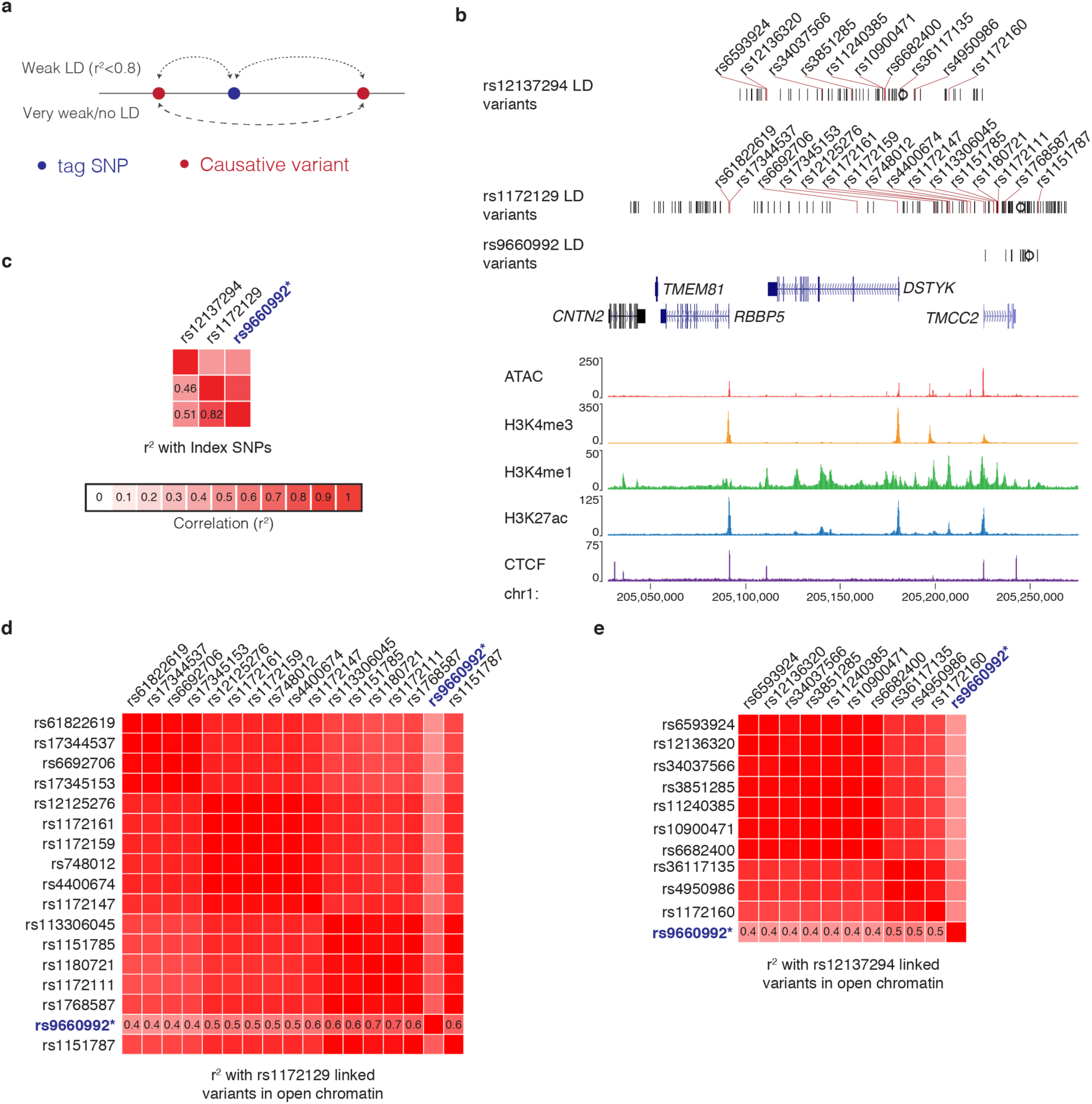
rs9660992 may be a tag SNP at the TMCC2 locus. **a,** Model for tag SNPs, whereby an index SNP may be identified through weak linkage disequilibrium (LD) to multiple unlinked causative variants. **b**, Map of RBC trait associated variants at the TMCC2 locus, with ATAC-seq and ChIP-seq chromatin marks from erythroid cells. Variants in open chromatin are red and labelled, index SNPs are marked with a circle. TMCC2 is a transmembrane and coiled-coil domain protein required for terminal erythropoiesis. **c**, Linkage analysis between index SNPs at the TMCC2 locus, the putative tag SNP rs9660992 is in blue. Linkage analysis between rs9660992 and variants in open chromatin that are in strong LD (r^2^≥0.8) with rs1172129 (**d**) and rs12137294 (**e**).

**Supplementary Fig. 13.**
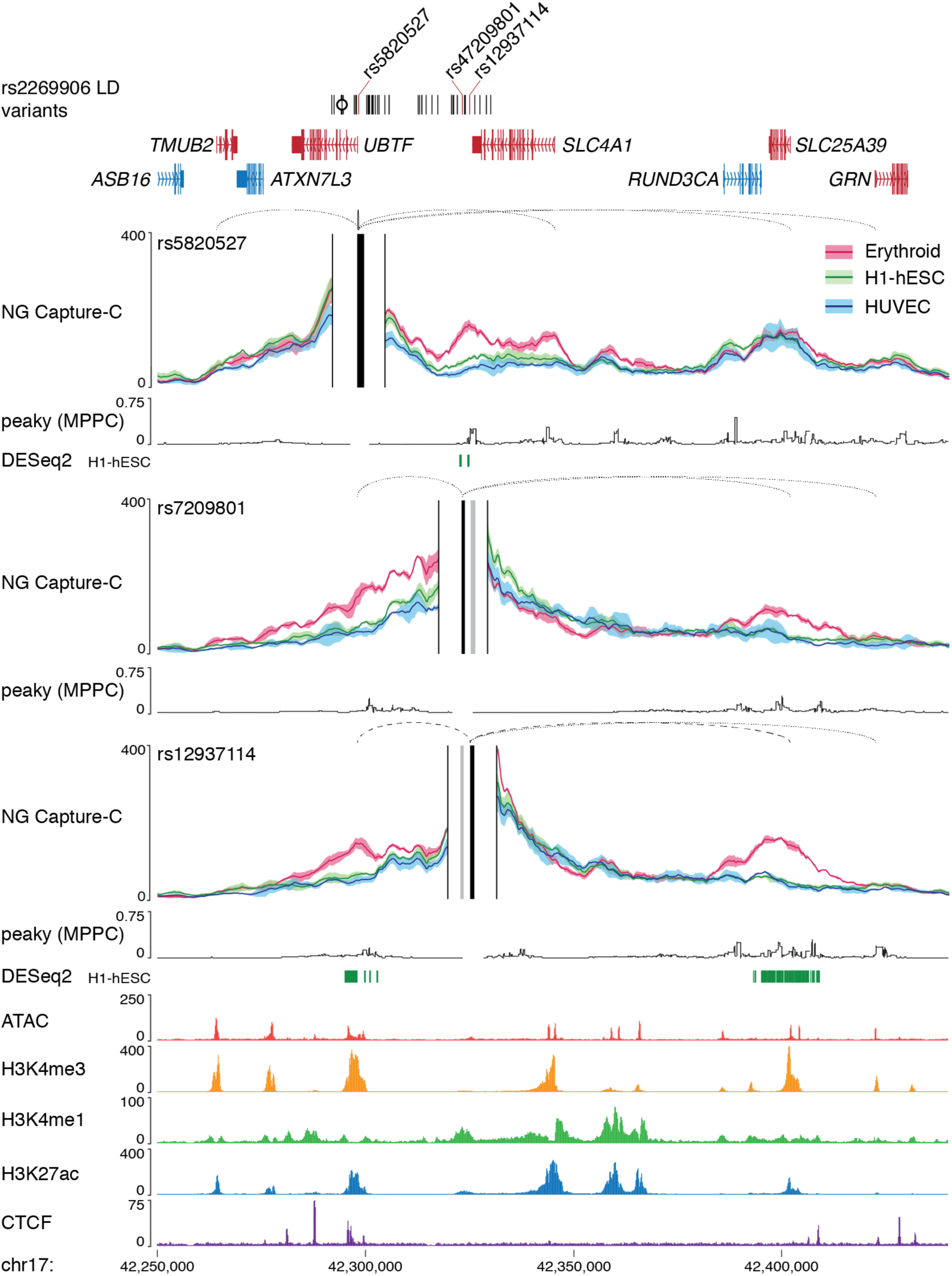
3C interactions at the Band 3 locus. NG Capture-C interaction profiles for three open chromatin intersecting variants at the SLC4A1 (which encodes Band 3) locus performed in erythroid, embryonic stem (H1-hESC) and umbilical vein endothelial (HUVEC) cells (n=3). Capture viewpoints and proximity exclusion regions (solid vertical lines) were designed for each open chromatin region and profiles show mean interactions (solid line) with one standard deviation (shading). Peaky values depict the Marginal Posterior Probability of Contact (MPPC) in erythroid cells. Promoter-variantsP interactions were identified by promoter proximity (solid loops), erythroid specific interactions (dashed loops; DESeq2 q-value < 0.05 shown as bars), and Bayesian modelling (dotted loops; peaky MPPC > 0.01). SNPs within open chromatin are red, as are variant interacting genes, the index SNP is marked with a circle. FPKM normalised ATAC-seq and ChIP-seq tracks are from erythroid cells. Key interactions were seen with both *SLC4A1*, an erythrocyte membrane anion transport protein and, and *SLC25A39* is a mitochondrial iron transport protein, both of which are required for erythropoiesis.

**Supplementary Fig. 14.**
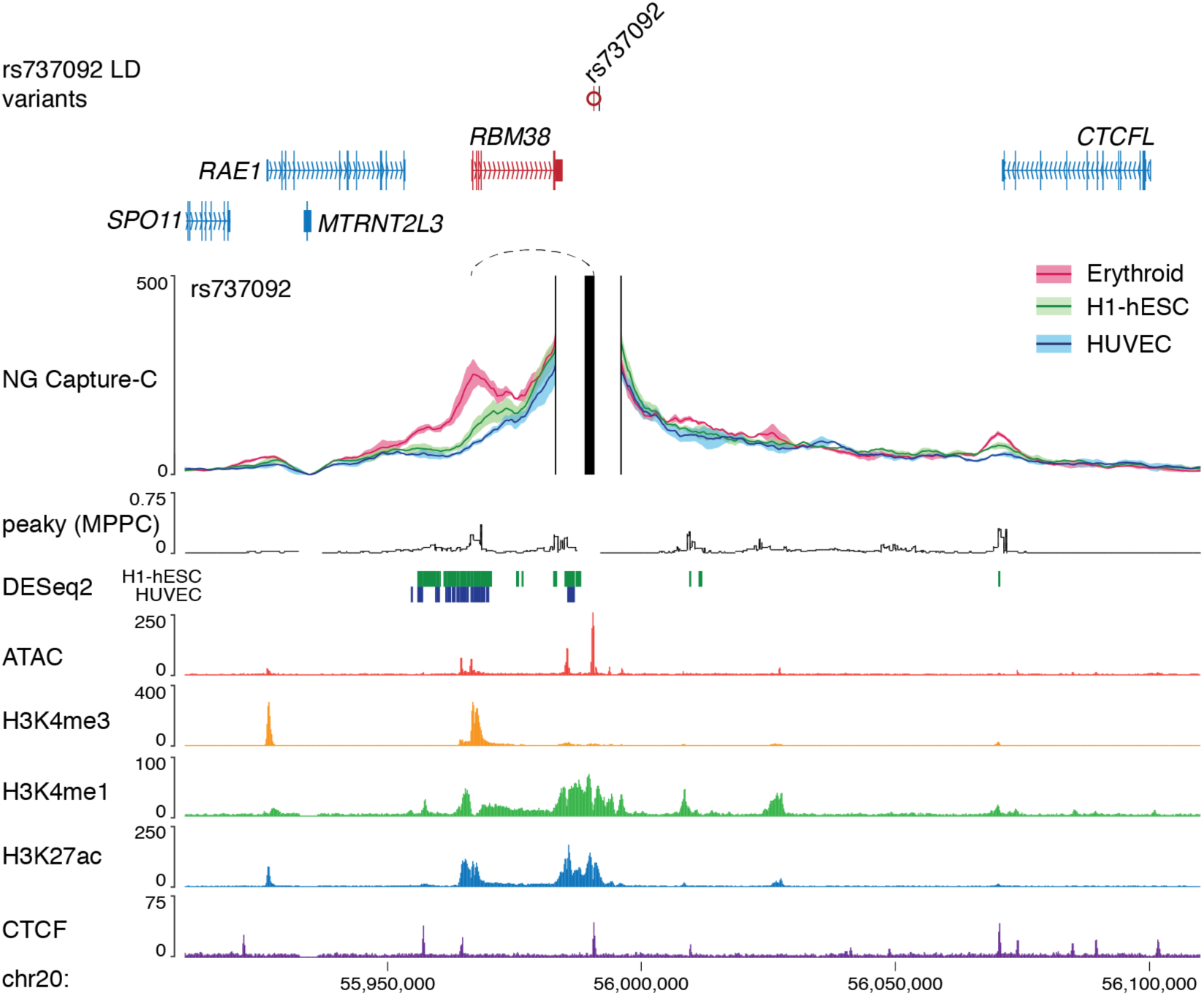
3C interaction profile for RNPC1 locus. 3C interaction profile for rs737092 from erythroid, embryonic stem (H1-hESC) and umbilical vein endothelial (HUVEC) cells (n=3). Capture viewpoints and proximity exclusion regions (solid vertical lines) were designed for each open chromatin regions and profiles show mean interactions (solid line) with one standard deviation (shading). Peaky values depict the MPPC in erythroid cells. Promoter-variant interactions were identified as erythroid-specific interactions (dashed loops; DESeq2 q-value < 0.05 shown as bars). Variants within open chromatin are red, as are variant interacting genes, the index SNP is marked with a circle. FPKM normalised ATAC-seq and ChIP-seq tracks are from erythroid cells. Interaction was seen with *RBM38*, which encodes the erythroid splicing regulator RNPC1.

**Supplementary Fig. 15.**
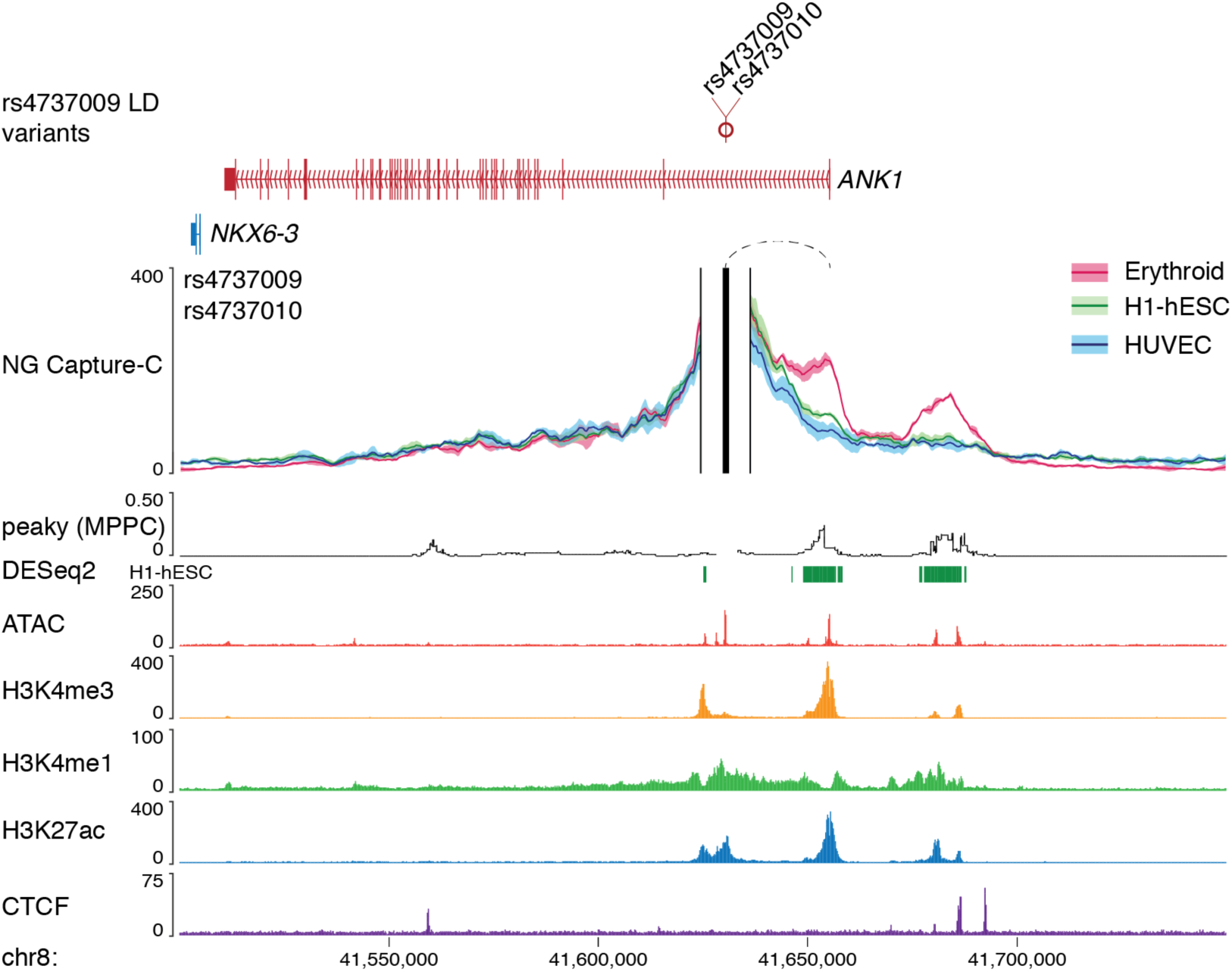
3C interaction profile for the Ankyrin locus. 3C interaction profile for rs4737009 and rs4737010 in erythroid, embryonic stem (H1-hESC) and umbilical vein endothelial (HUVEC) cells (n=3). Capture viewpoints and proximity exclusion regions (solid vertical lines) were designed for each open chromatin regions and profiles show mean interactions (solid line) with one standard deviation (shading). Peaky values depict the MPPC in erythroid cells. Promoter-variant interactions were identified as erythroid specific (dashed loops; DESeq2 q-value < 0.05 shows as bars). Variants within open chromatin are red, as are variant interacting genes, the index SNP is marked with a circle. FPKM normalised ATAC-seq and ChIP-seq tracks are from erythroid cells. Interaction was seen with *ANK1*, which encodes Ankyrin-1 – an integral erythroid membrane protein often missing in hereditary spherocytosis patients.

**Supplementary Fig. 16.**
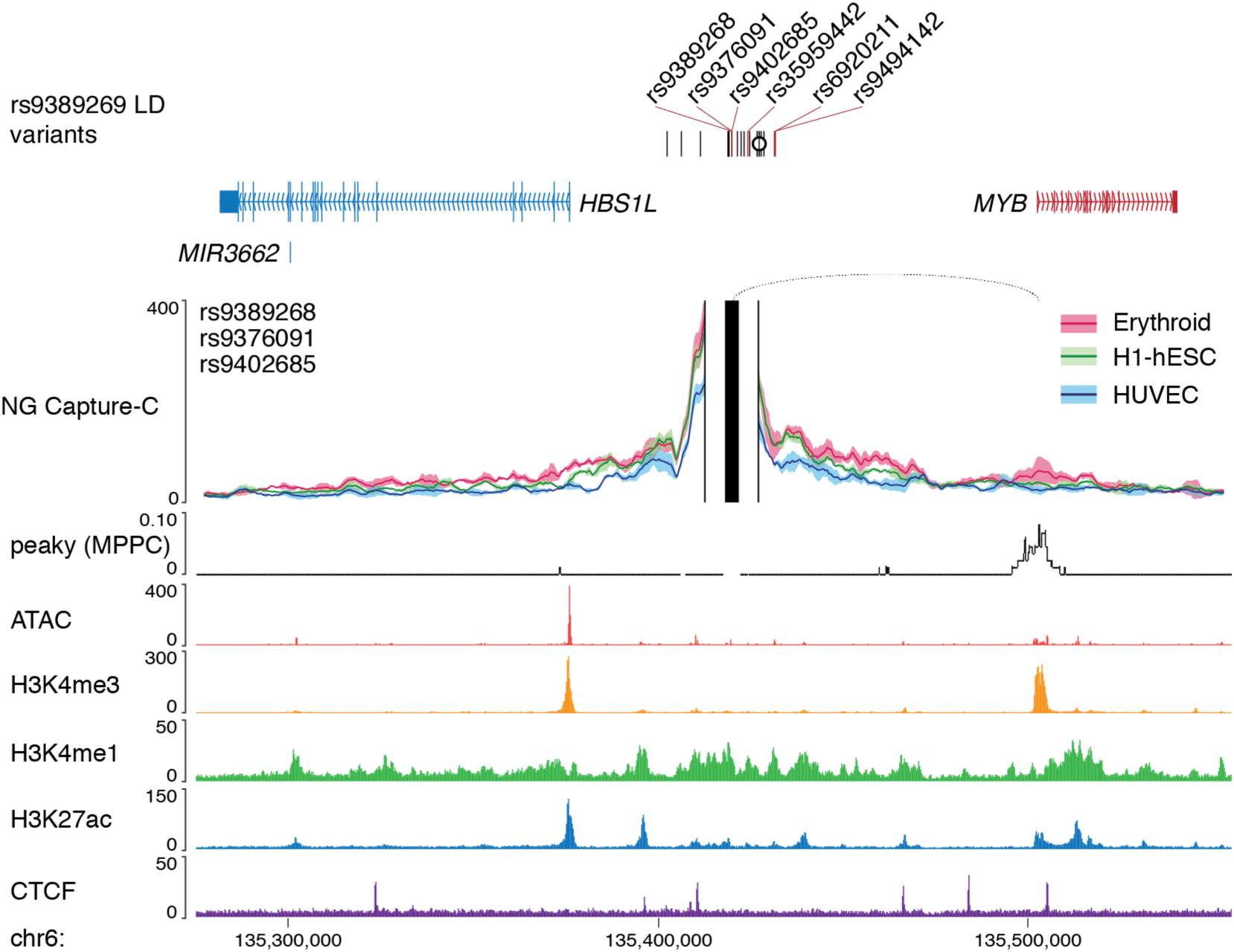
3C interaction profile for the c-Myb locus. 3C interaction profile for rs9389268, rs9376091, and rs9402685 in erythroid, embryonic stem (H1-hESC) and umbilical vein endothelial (HUVEC) cells (n=3). Capture viewpoints and proximity exclusion regions (solid vertical lines) were designed for each open chromatin regions and profiles show mean interactions (solid line) with one standard deviation (shading). Peaky values depict the MPPC in erythroid cells. Interaction with *MYB* was detected by peaky (dotted loop; MPPC > 0.01). Variants within open chromatin are red, as are variant interacting genes, the index SNP is marked with a circle. FPKM normalised ATAC-seq and ChIP-seq tracks are from erythroid cells. Interaction was seen with the *MYB*, which encodes c-Myb a key regulatory transcription factor involved in erythroid progenitor proliferation and differentiation.

**Supplementary Fig. 17.**
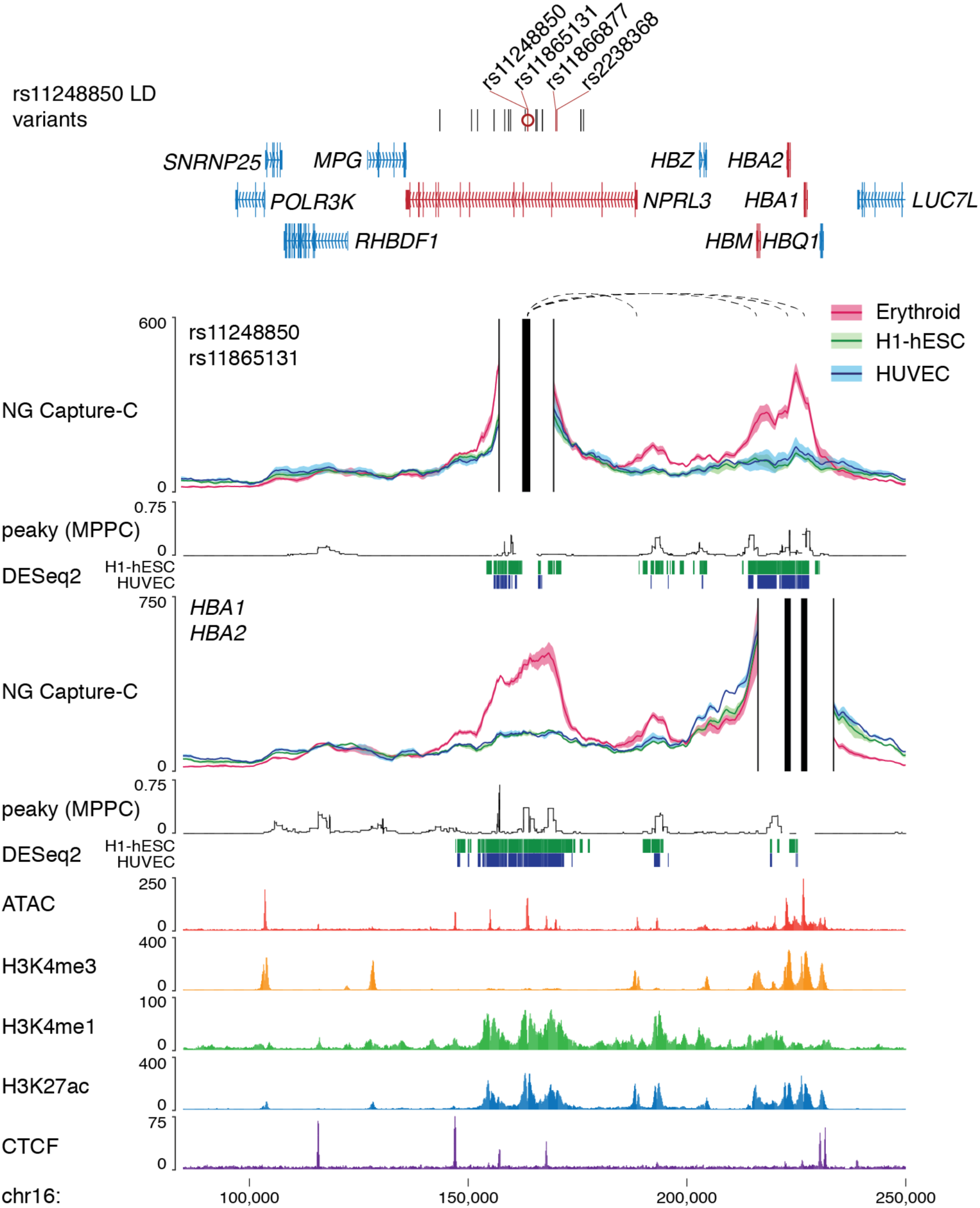
3C interaction profile for the α-globin locus. 3C interaction profile for rs11248850 and rs11865131, which lie in a conserved HBA1 and HBA2 enhancer, as well as reciprocal promoter capture in erythroid, embryonic stem (H1-hESC) and umbilical vein endothelial (HUVEC) cells (n=3). Capture viewpoints and proximity exclusion regions (solid vertical lines) were designed for each open chromatin regions and profiles show mean interactions (solid line) with one standard deviation (shading). Peaky values depict the MPPC in erythroid cells. Promoter-variant interactions were identified as erythroid specific (dashed loops; DESeq2 q-value < 0.05 show as bars). Variants within open chromatin are red, as are variant interacting genes, the index SNP is marked with a circle. FPKM normalised ATAC-seq and ChIP-seq tracks are from erythroid cells.

**Supplementary Fig. 18.**
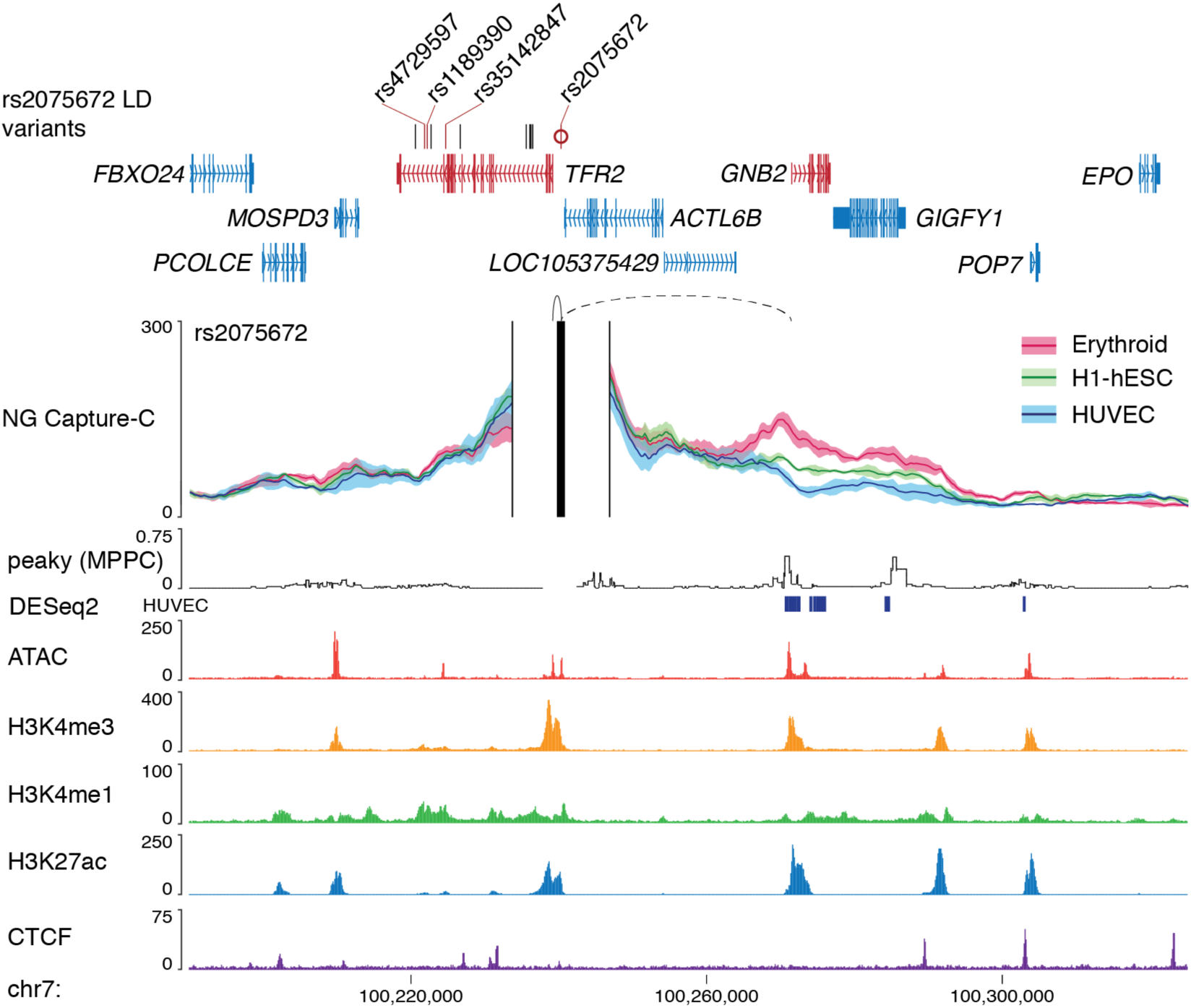
3C interaction profile for the TfR2 locus. 3C interaction profile for rs2075672 in erythroid, embryonic stem (H1-hESC) and umbilical vein endothelial (HUVEC) cells (n=3). Capture viewpoints and proximity exclusion regions (solid vertical lines) were designed for each open chromatin regions and profiles show mean interactions (solid line) with one standard deviation (shading). Peaky values depict the MPPC in erythroid cells. Promoter-variant interactions were identified by promoter proximity (solid loops) and erythroid specific interactions (dashed loops; DESeq2 q-value < 0.05 shows as bars). Variants within open chromatin are red, as are variant interacting genes, the index SNP is are marked with a circle. FPKM normalised ATAC-seq and ChIP-seq tracks are from erythroid cells. Interaction was seen with *TFR2*, which encodes the erythroid and liver hepcidin receptor Transferrin receptor 2 (TfR2), and *GNB2*, which encodes the signalling G-protein subunit B.

**Supplementary Fig. 19.**
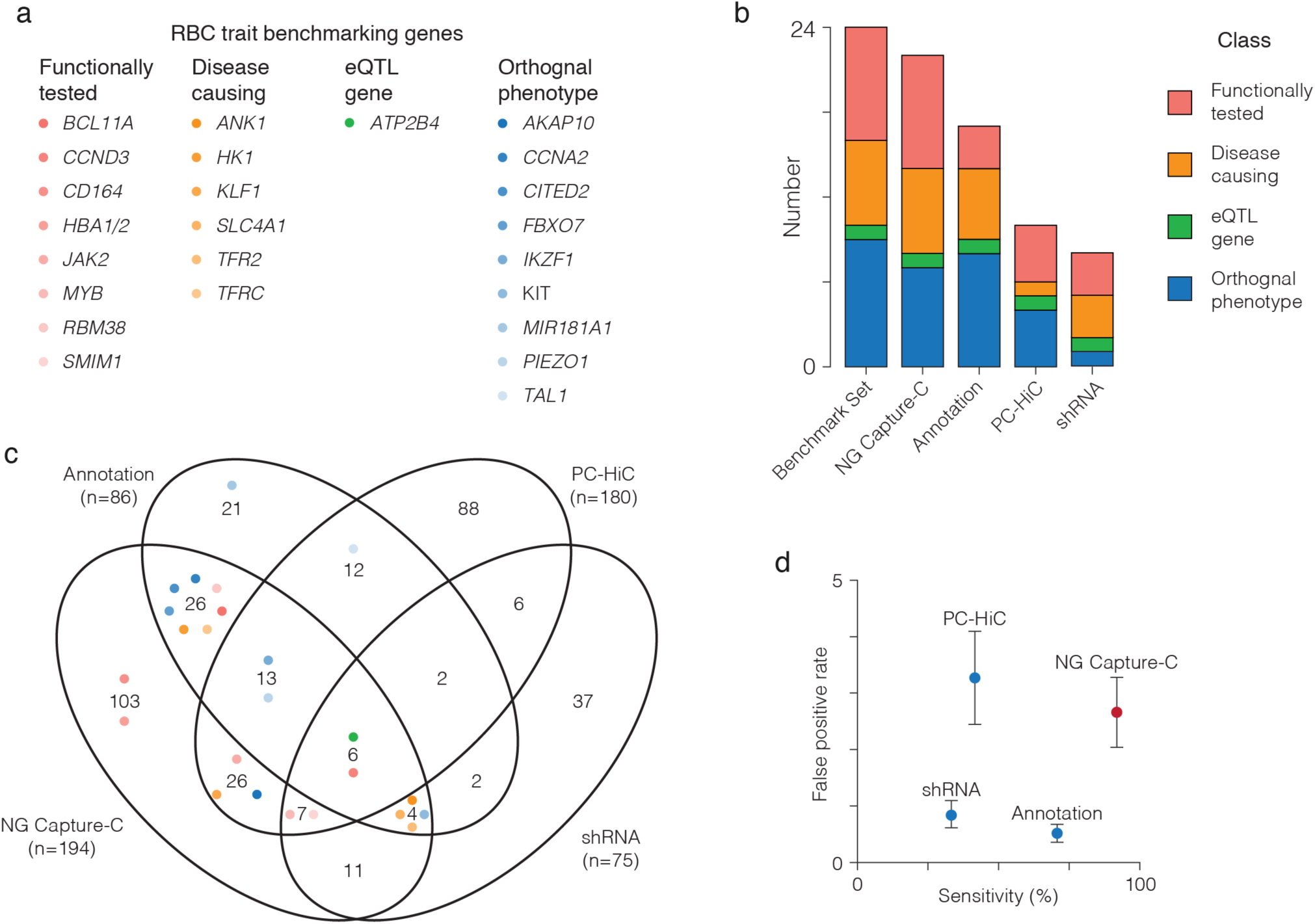
Comparison of methods for identification of GWAS effector genes. **a,** Classification of 24 most-likely effector genes used for benchmarking as identified through known function in erythropoiesis. **b,** Number of benchmarking genes identified by four different methods including NG Capture-C, Promoter Capture-HiC (PC-HiC), a gene centric shRNA screen, and an annotation-based method utilising coding variants, eQTLS, nearby genes and gene relationships among implicated loci (GRAIL). **c,** Venn comparison of all genes identified by the four different methods across the 53 NG Capture-C targeted chromosome regions with dots for each of the benchmarking genes. **d**, Percent of 24 benchmarking genes identified (sensitivity) and average number of false positives per locus (false positive rate) for each of the four methods. Error bars show standard error of the mean (n = 24).

**Supplementary Fig. 20.**
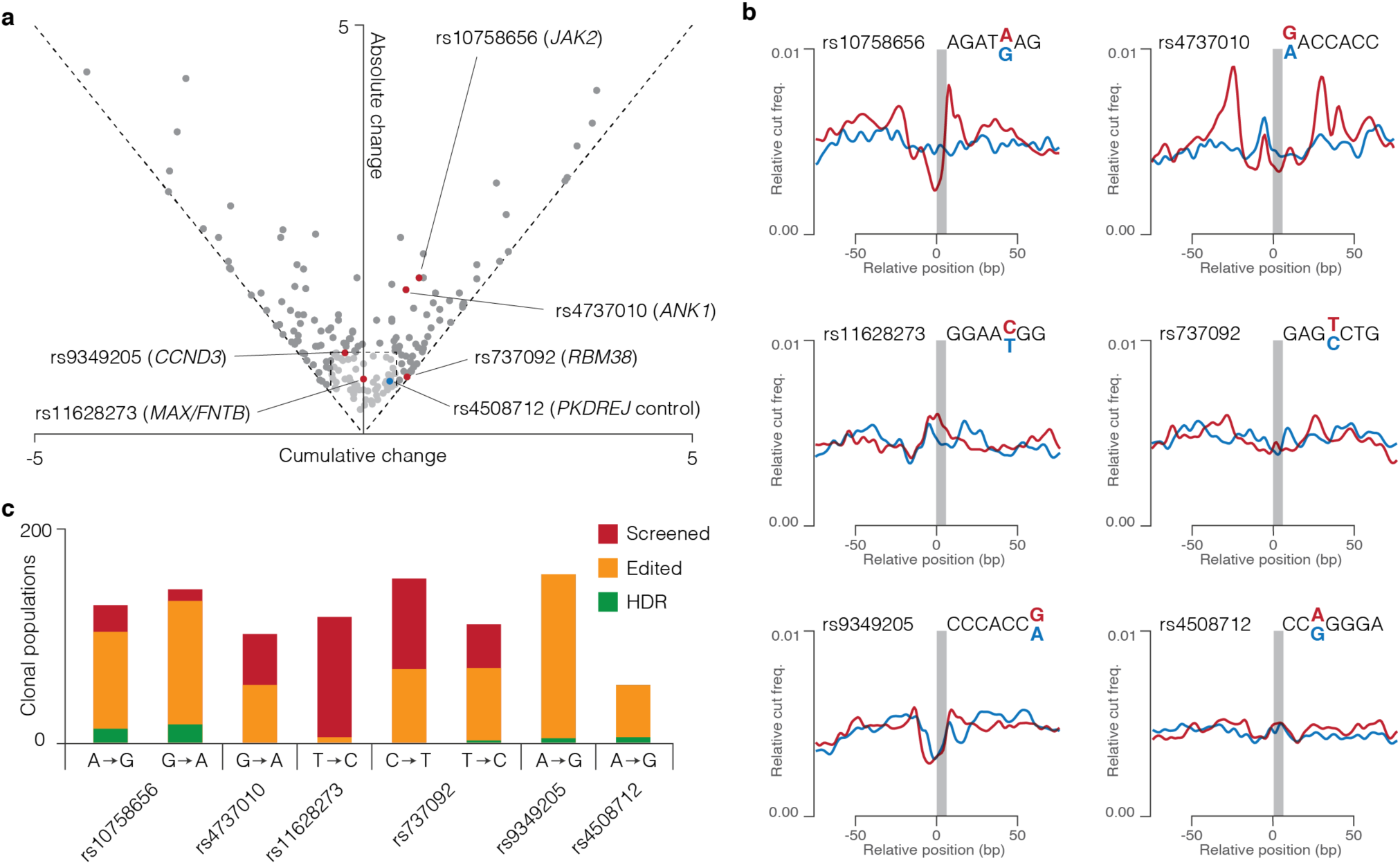
Pipeline prioritized SNPs were edited by CRISPR/Cas9. **a,** Five SNPs with candidate effector genes (red dots) and one control SNP (blue dot) were selected for *in vivo* modelling. **b,** Selected SNPs showed a range of predicted Sasquatch effects and alleles were directly modelled by CRISPR/Cas9 facilitated homology directed repair (HDR). **c,** Clonal HUDEP-2 populations were derived by editing and single-cell sorting and screened by next generation sequencing to detect changes. Bar height shows total number of clonal populations screened, with variable rates of editing (indels and HDR, yellow) and specific changes (HDR, green) observed across the loci.

**Supplementary Fig. 21.**
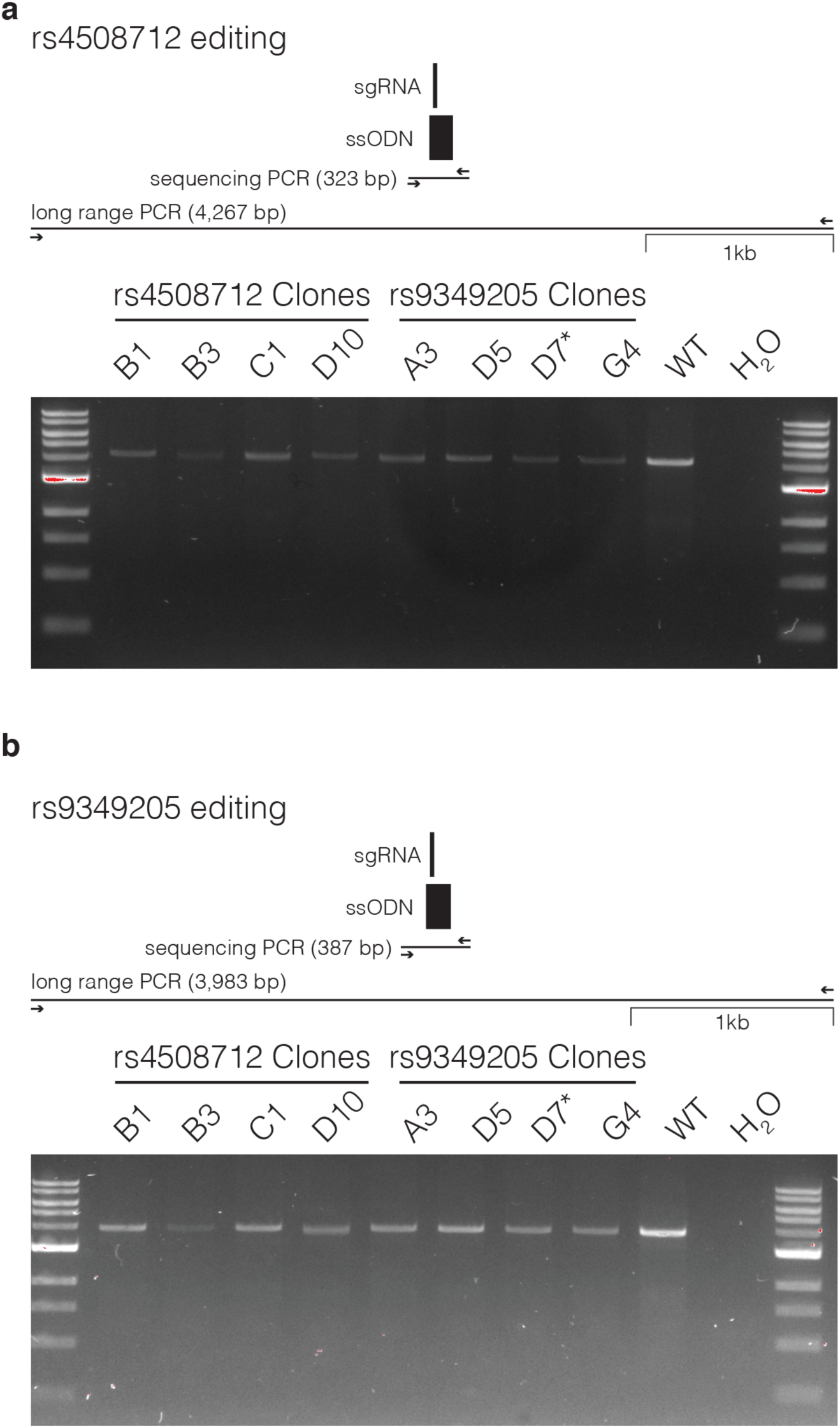
Long range PCR to detect loss of heterozygosity. Long range PCR was carried out for the both rs4508712 (**a**) and rs9349205 (**b**) loci on clones identified by sequencing as homozygous for the desired editing events. *Re-sequencing identified a 2bp deletion leading to the exclusion of this clone.

**Supplementary Fig. 22.**
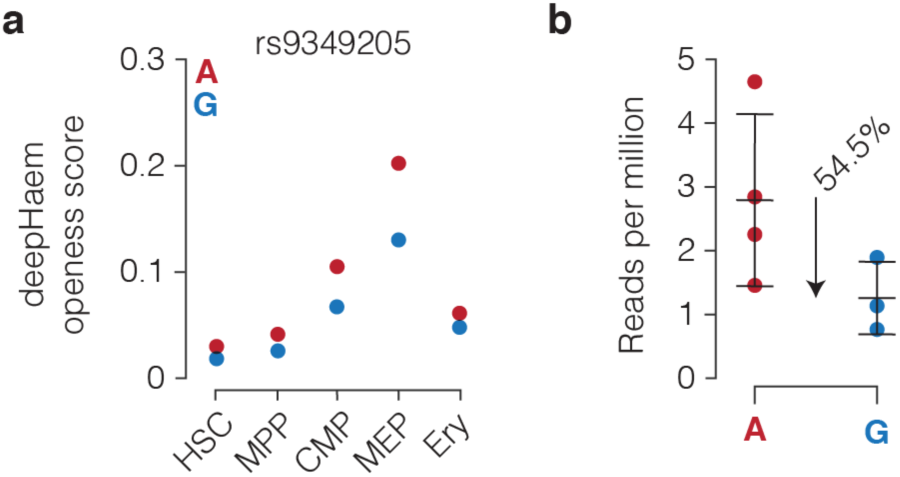
deepHaem openness score for rs9349205. **a,** DeepHaem uses a deep convolutional neural network machine learning approach to predict the likelihood of chromatin openness (or “openness score”) ranging from 0-1. These prediction scores were generated through training on a binary yes or no for openness, but do have a positive correlation with observed levels ATAC-seq signal (see Supplementary Figure 7). For this study, differences between variant alleles greater than 0.1 (or 10% of the maximum openness score) were classed as damaging. For rs9349205 a slight difference between the two alleles is seen with deepHaem, though this does no cross the strict threshold for classification as damaging. **b,** Count of ATAC-seq reads per million mapped reads in the rs9349205 containing peak (chr6:41,924,682-41,925,483, hg19) in undifferentiated human umbilical derived erythroid progenitors (HUDEP-2) cells. Each circle is an independently isolated clone homozygous for either A or G at rs9349205, bars show mean and one standard deviation.

**Supplementary Fig. 23.**
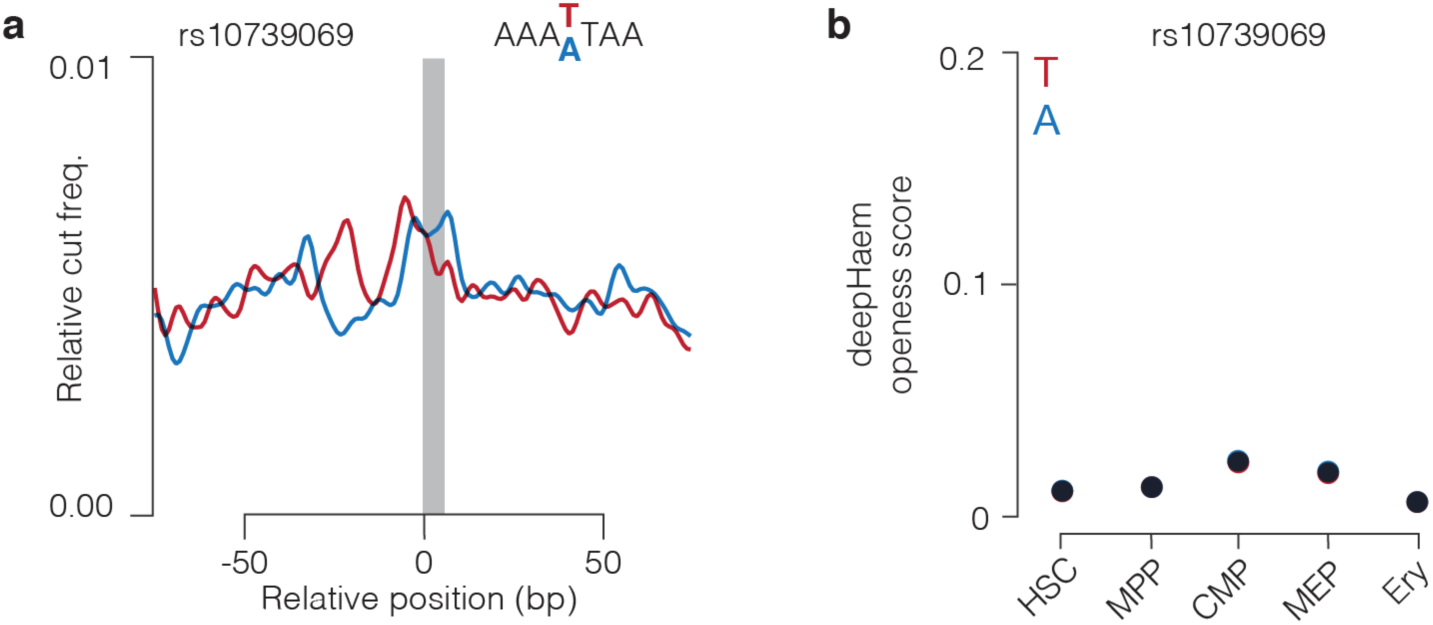
rs10739069 is not predicted to be damaging. rs10739069 was found within an open chromatin region, however the Sasquatch footprint (**a**) and deepHaem openness scores (**b**) show it is unlikely to be a damaging RBC trait SNP.

**Supplementary Fig. 24.**
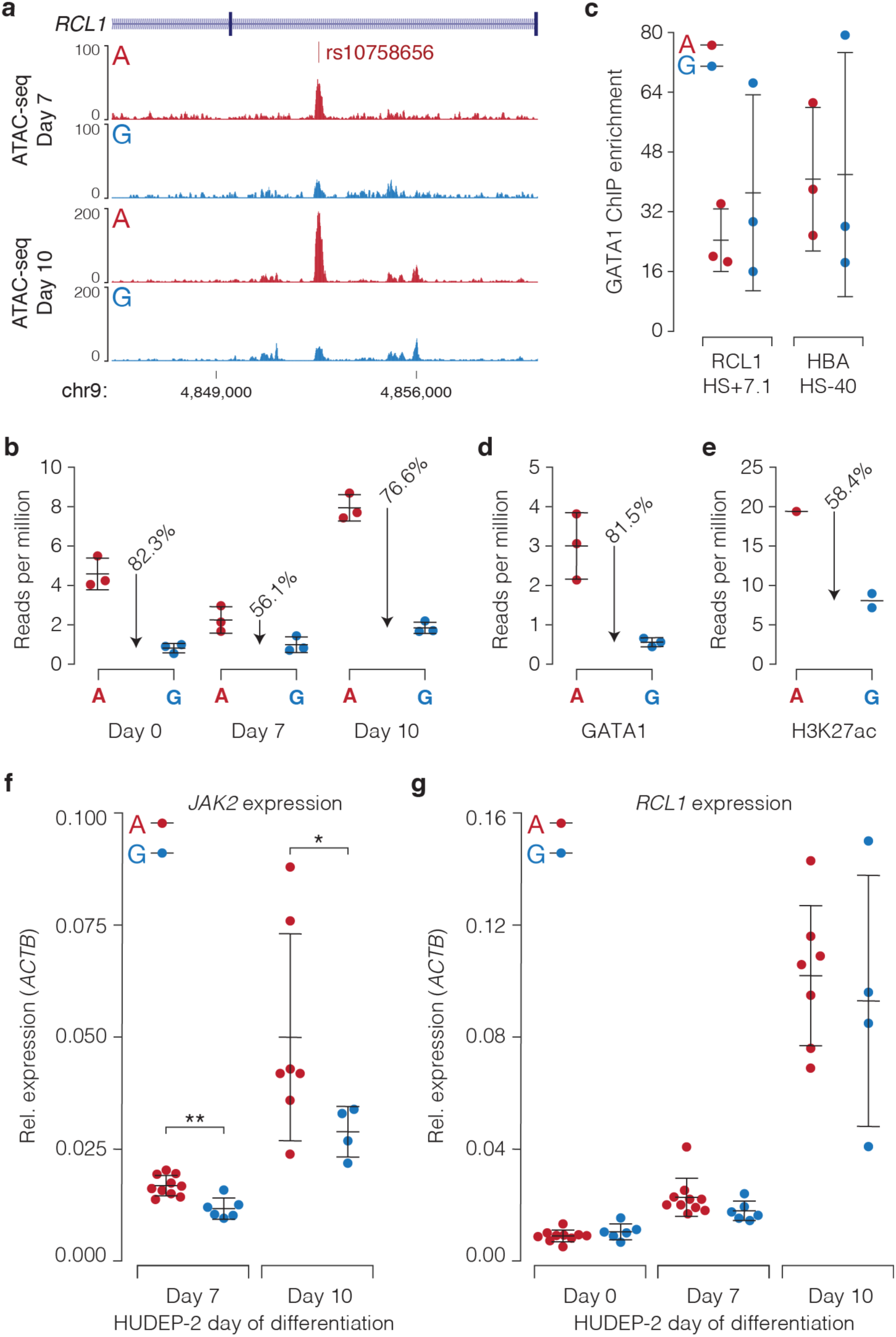
rs10758656 is specific for regulation of *JAK2*. **a,** FPKM normalised ATAC-seq (n=3) from differentiating HUDEP-2 clones homozygous for either rs10758656 allele. **b,** Count of ATAC-seq reads per million mapped reads in the rs10758656 containing peak (chr9:4,852,237-4,853,036, hg19) in undifferentiated human umbilical derived erythroid progenitors (HUDEP-2) cells. **c,** Real-time quantitative PCR for GATA1 ChIP enrichment at two control open chromatin regions, one upstream of rs10758656 near the RCL1 promoter (chr9:4799898-4799984, hg19) and a well characterized *HBA1/2* enhancer (HS-40; chr16:163615-163730, hg19) in rs10758656^A/A^ and rs10758656^G/G^ clones. Enrichment was calculated using the ΔΔCt method against a background region. Count of reads per million mapped reads in the rs10758656 containing peak from ChIP-seq of GATA1 (**d**) and H3K27ac (**e**) at day 7 of differentiation. Real time reverse-transcriptase PCR of *JAK2* (**f**) and the gene nearest to rs10758656, *RCL1* (**g**) *–* which encodes RNA terminal phosphate cyclase-like 1 – in differentiating HUDEP-2 clones edited to be homozygous for both alleles of rs10758656 (n≥4). Mann-Witney test, **p=0.0030, *p=0.0364. For all graphs each circle is an independently isolated clone homozygous for either A or G at rs10758656, bars show mean and one standard deviation.

**Supplementary Fig. 25.**
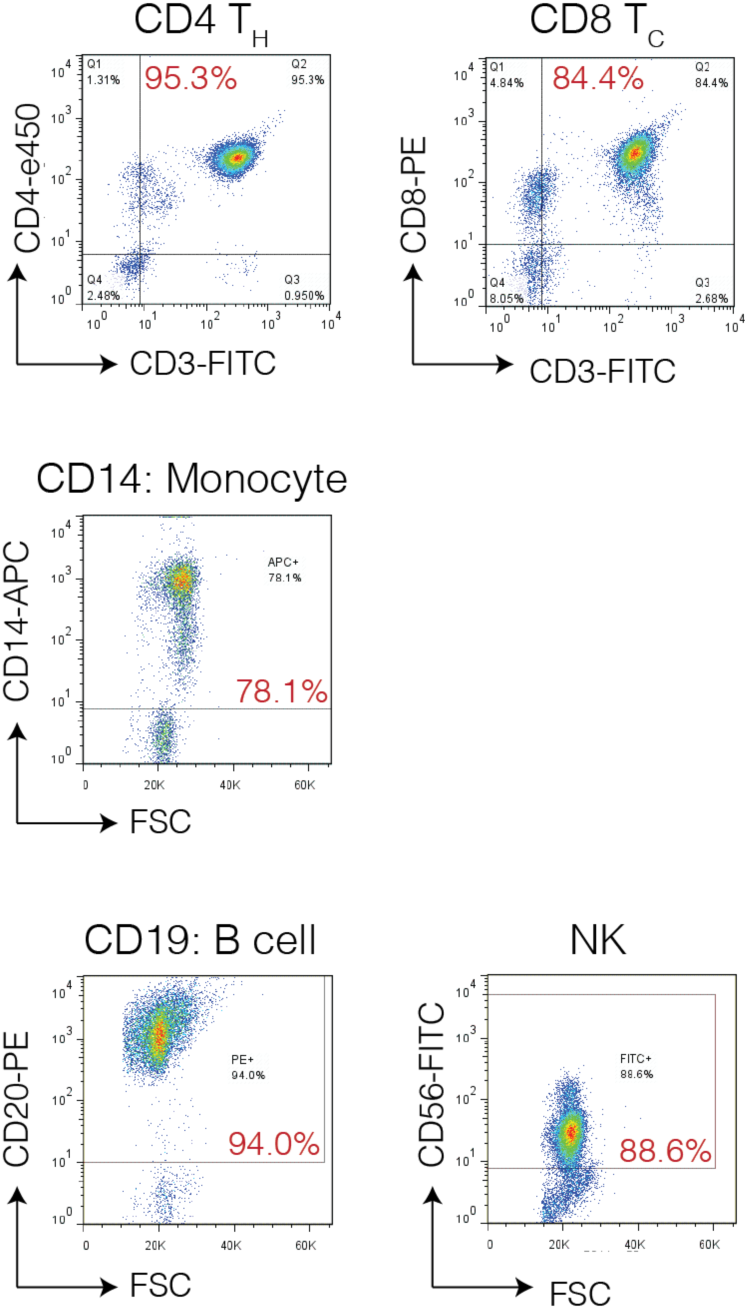
Population purities for MACS isolated immune cell populations. MACS sorted populations were stained with a specific cocktail of cell markers to determine purity. Gates were set using a negative staining cell population.

**Supplementary Table 1. RBC Prioritized variants.**

a. Overview of number of variants analysed by the integrated platform
b. List of prioritized variants with supporting evidence and gene interactions.
c. List of previously validated variants.
d. List of benchmarking RBC trait genes.

**Supplementary Table 2. Growth media recipes.**

a. CD34 Differentiation.
b. HUDEP-2 Differentiation.

**Supplementary Table 3. Oligonucleotides used for genome editing.**

a. Genotyping primers.
b. Guide RNA.
c. ssODN.
d. Long range PCR primers.

**Supplementary Table 4. List of antibodies.**

a. Flow Cytometry.
b. Chromatin Immunoprecipitation.

**Supplementary Table 5. Quantitative PCR primers**

a. TaqMan Expression.
b. DpnII 3C library digestion.
c. ChIP Enrichment.

**Supplementary Table 6. Sample Sequencing Depth.**

a. RNA-seq.
b. NG Capture-C.
c. ATAC-seq.
d. ChIP-seq.

**Supplementary Table 7. List of biotinylated NG Capture-C oligonucleotides.**

a. Enhancer Oligonucleotides.
b. Promoter Oligonucleotides.

**Supplementary Table 8. Bed files of open chromatin peak calls.**

a. HSC.
b. MPP.
c. CMP.
d. MEP.
e. Erythroid Day 7.
f. Erythroid Day 10.
g. Erythroid Day 13.
h. Erythroid Day 17.
i. Erythropoiesis Staging.
j. H1-hESC.
k. HUVEC.
l. CD4.
m. CD8.
n. CD14.
o. CD19.
p. Natural Killer.

